# Voice Patterns in Schizophrenia: A systematic Review and Bayesian Meta-Analysis

**DOI:** 10.1101/583815

**Authors:** Parola Alberto, Simonsen Arndis, Bliksted Vibeke, Fusaroli Riccardo

## Abstract

Voice atypicalities have been a characteristic feature of schizophrenia since its first definitions. They are often associated with core negative symptoms such as flat affect and alogia, and with the social impairments seen in the disorder. This suggests that voice atypicalities may represent a marker of clinical features and social functioning in schizophrenia. We systematically reviewed and meta-analyzed the evidence for distinctive acoustic patterns in schizophrenia, as well as their relation to clinical features. We identified 46 articles, including 55 studies with a total of 1254 patients with schizophrenia and 699 healthy controls. Summary effect sizes (Hedges’g and Pearson’s r) estimates were calculated using multilevel Bayesian modeling. We identified weak atypicalities in pitch variability (*g* = - 0.55) related to flat affect, and stronger atypicalities in proportion of spoken time, speech rate, and pauses (*g*’s between -0.75 and -1.89) related to alogia and flat affect. However, the effects were mostly modest (with the important exception of pause duration) compared to perceptual and clinical judgments, and characterized by large heterogeneity between studies. Moderator analyses revealed that tasks with a more demanding cognitive and social component showed larger effects both in contrasting patients and controls and in assessing symptomatology. In conclusion, studies of acoustic patterns are a promising but, yet unsystematic avenue for establishing markers of schizophrenia. We outline recommendations towards more cumulative, open, and theory-driven research.

## Introduction

Individuals with schizophrenia display atypical voice patterns, qualitatively described in terms of poverty of speech, increased pauses, distinctive tone and intensity of voice (Alpert et al., 2000; Andreasen et al., 1985; Cohen et al., 2016, 2012b; Galynker et al., 2000; Hoekert et al., 2007; Trémeau et al., 2005). Voice atypicalities have been reported since the first definitions of the disorder (Bleuler, 1911; Kraepelin, 1919), are used in the clinical assessment process, and assume an even stronger relevance in the light of growing findings associating voice patterns to cognitive function, emotional states, and social engagement (Cohen and Elvevåg, 2014; Cohen and Hong, 2011; Faurholt-Jepsen et al., 2016; Nevler et al., 2017; Pisanski et al., 2016; Trigeorgis et al., 2016; Tsanas et al., 2011; Wang et al., 2015; Williams et al., 2014; Yin et al., 2007).

Voice atypicalities may thus constitute a window into the underlying clinical and cognitive features of the disorder. Indeed, they have been associated with core negative symptoms of schizophrenia such as blunted affect (e.g. diminished emotional expression, lack of vocal intonation), and alogia (e.g. poverty of speech, latency of speech and blocking) (Murray Alpert et al., 2000; Andreasen, 1984; Cohen et al., 2012; Millan, Fone, Steckler, & Horan, 2014; Ross, Orbelo, Cartwright, et al., 2001). Negative symptoms are included among the primary diagnostic criteria of schizophrenia (DSM-V), and are associated with early age of onset, poor social and functional outcome, reduced quality of life, and poor response to medication and treatment (Couture et al., 2011; Häfner et al., 1999; Rabinowitz et al., 2012; Tandon et al., 2008). Vocal expression also reflects a key component of social communication, a domain frequently impaired in individuals with schizophrenia (Bambini et al., 2016; Bosco et al., 2019; Brüne and Bodenstein, 2005; Champagne-Lavau and Stip, 2010; Colle et al., 2013; Meilijson et al., 2004; Parola et al., 2018). Difficulties in controlling voice to express affective states, e.g. emotional, mood or motivational states, or to mark relevant information may dramatically reduce the ability of these individuals to communicate effectively in social context. Impairments in social communication may in turn lead to experience of failure in social situations, and atypical voice qualities may generate more negative attitudes and social judgments on the part of others than toward speakers with typical voice qualities (e.g. Altenberg & Ferrand, 2006). This, in turn, may result in social withdrawal and the further aggravation of social cognitive impairments (Bambini et al., 2016; Bowie and Harvey, 2008; Del-Monte et al., 2013; Sparks et al., 2010; Tan et al., 2014; Thoma et al., 2009). Voice atypicalities may thus represent an important marker that parallels both clinical features and social cognitive functioning of individuals with schizophrenia over time (Cohen et al., 2016; Rapcan et al., 2010; Tahir et al., 2019).

By understanding the implications of voice abnormalities, we can provide a first step to develop more effective tools to assist clinicians in assessing this heterogeneous disorder. For example, different speech and voice features such as pause number and duration, speech rate and pitch and intensity variability, have been proved to be potential indicators of cognitive load both in healthy individuals (Berthold and Jameson, 1999; Cohen et al., 2015; Khawaja et al., 2008; Yin et al., 2007), and individual with schizotypal disorder (Cohen et al., 2012a). In addition, voice analysis may potentially allow to assess the response to psychosocial or pharmacological treatment over longer periods using objective and quantitative indices, and enhance the capability of clinicians to capture the complex relationship between emotion regulation, expressive behavior, social perception and cognitive and clinical features of the disorder (e.g. Ben-Zeev et al., 2017; Dahlgren et al., 2018; Tahir et al., 2019) . Finally, vocal abnormalities may be related to neuromotor disorders frequently associated with schizophrenia and its neurodevelopmental pathophysiology (Cannizzaro et al., 2005; Konopka and Roberts, 2016; Matsumoto et al., 2013; Walther, 2015; Walther and Strik, 2012), or secondary induced by antipsychotic medications (Peluso et al., 2012; Tenback et al., 2010). Clarifying the vocal abnormalities associated with schizophrenia may thus provide tools to support the assessment of quantitative measures of cognitive and clinical features related to voice, such as cognitive load, negative symptomatology and affective states, but also contribute toward a better understanding of the nature and pathophysiology of the disorder.

Despite the importance of studying vocal expression in schizophrenia, and the routine assessments performed using interview-based clinical rating scales, our understanding of voice abnormalities in schizophrenia is limited. Previous work on voice atypicalities can be organized into three categories: qualitative perceptual ratings, quantitative acoustic analyses, and multivariate machine learning (ML) investigations. Most previous studies employing qualitative ratings reported robust differences between patients with schizophrenia and healthy controls (HC) across several perceptual features of their voice, often described as lack of inflection, poverty of speech and latency of speech (Cohen et al., 2014; Emmerson et al., 2009; Hoekert et al., 2007; Mueser et al., 1994). However informative, qualitative rating scales have serious limitations. They rely on raters’ expertise and intuition, thus lacking scalability to large corpora, that is, they are expensive and time-consuming to apply to a large number of recordings and require specialized training of the clinicians. Further, they display low sensitivity to complex and multivariate acoustic patterns and variations in context and time (Alpert, Pouget, & Silva, 1995; Cohen & Elvevåg, 2014; Cohen, Mitchell, Docherty, & Horan, 2016a; Cohen et al., 2012). A different approach involves the use of automated analysis of speech to identify acoustic features of vocal production, arguably with a greater reliability, sensitivity and validity. However, such studies have so far reported smaller and seemingly more contradictory findings: some indicate slower speech (Tavano et al., 2008), more pronounced pauses (Cannizzaro et al., 2005; Martínez-Sánchez et al., 2015; Rapcan et al., 2010) and reduced prosodic variability (Compton et al., 2018; Martínez-Sánchez et al., 2015; Ross et al., 2001); while others indicate no reliable acoustic differences between individuals with schizophrenia and HC (Cohen et al., 2016; Docherty, 2012; Meaux et al., 2018). A meta-analysis of 13 studies (Cohen et al., 2014) suggests large differences between individuals with schizophrenia and HC on pause and speech duration, and more modest on intensity and pitch variability. While informative, the number of studies included in the meta-analysis was small compared to the currently available literature and, given the high heterogeneity of patients with schizophrenia, a more systematic review accounting for the potential sources of heterogeneity in the effects is required: individual differences (e.g. gender, age and education), contextual factors (e.g. type of task) and clinical features (e.g. symptomatology and medication). This is crucial if we want to understand the mechanisms underlying voice atypicalities. Different mechanisms may lead to different acoustic patterns in different contexts, e.g. cognitive impairment leading to longer pauses, and even more so when the cognitive and social demands of the task are increased. On the contrary, voice atypicalities due to differences in fine motor control of the vocal folds should be similar across tasks. Further, a few studies have adopted a more fine-grained perspective, and assessed the relationship between acoustic measures and clinical features with some promise; however, the findings are still very sparse (Alpert, Shaw, Pouget, & Lim, 2002; Alpert et al., 2000; Cohen et al., 2016a; Meaux et al., 2018; Zhang, Pan, Gui, Zhu, & Cui, 2016).

Finally, more recent studies have tried to capitalize on the technological advancements in speech signal processing, and the application of multivariate ML techniques to better capture the complex, multivariate and often non-linear nature of acoustic patterns (Bone et al., 2017; Huys et al., 2016; for an introduction to ML techniques in the context of voice analysis see also the appendix to Fusaroli, Lambrechts, Bang, Bowler, & Gaigg, 2017). These studies extract more nuanced acoustic measures, e.g. spectral and glottal features, and assess how accurately the diagnosis can be identified only relying on acoustic measures. The results are promising (Bonneh, Levanon, Dean-Pardo, Lossos, & Adini, 2011; Faurholt-Jepsen et al., 2016; Martínez-Sánchez et al., 2015; Rapcan et al., 2010; Tsanas et al., 2011; Williams et al., 2014), but a complete and comparative overview of the findings in schizophrenia is currently missing. Crucially, the reliability of ML results has been shown to be strongly dependent on the availability of large datasets and the validation of the findings across datasets (Bone et al., 2016; Chekroud, 2018; Foody, 2017; James et al., 2013; Van Der Ploeg et al., 2014), which presence we wanted to assess in the literature on voice in schizophrenia.

Despite the promise of acoustic markers of clinical features in schizophrenia, it is yet unclear how to quantify them, that is, which acoustic features we should focus on, and the evidence for their relation to specific clinical features of the disorder. The aim of the present study was to fill this gap by systematically reviewing and meta-analyzing the current state of evidence for acoustic atypicalities in schizophrenia as a whole as well as their relation to the specific clinical features. Further, we evaluated the size and availability of previous datasets, and the attitudes towards data sharing of the authors of the studies reviewed to assess whether a more cumulative science of voice atypicalities in schizophrenia can be attempted. Note that the aim of this meta-analysis is less to provide a more accurate estimation of the voice atypicalities in schizophrenia than it is to provide the basis for more effective future studies, by identifying current practices, issues and promising venues.

## Methods

### Inclusion criteria for literature search

We adopted the Preferred Reporting Items for Systematic Reviews and Meta-Analyses Guidelines (PRISMA, Stewart et al., 2015) for transparent reporting of a systematic review. We pre-registered our protocol by specifying a priori the study rationale, eligibility criteria, search strategy, moderator variables, and statistical analyses (see https://bit.ly/2EEFeQZ). The literature search was conducted on Pubmed and Google Scholar, the latter including dissertations and unpublished manuscripts. The search terms used were (prosody OR inflection OR intensity OR pitch OR fundamental frequency OR speech rate OR voice quality OR acoustic OR intonation OR vocal) AND (schizo*). The search was conducted on August 21 2017, and updated on April 12 2018. We complemented the list by performing a backward and forward literature search: we screened the bibliography of the papers found, and the papers citing them as identified by Google Scholar.

Articles were screened for eligibility by two authors (AP and AS). Study selection was conducted according to the following inclusion criteria: (a) empirical study, (b) quantification of acoustic features in the vocal production of participants with schizophrenia or schizoaffective disorder^1^ (c) sample including at least two individuals with schizophrenia or schizoaffective disorder, (d) inclusion of a non-clinical comparison group, or an assessment of variation in acoustic features in relation to severity of clinical features. Clinical comparison groups (e.g. with depression) were excluded because the limited number of studies did not permit meta-analytic estimations. Fig. 1 shows the flow-diagram of study selection. We report the assessment of the risk of bias in the Supplementary Materials.

**Figure 1.**
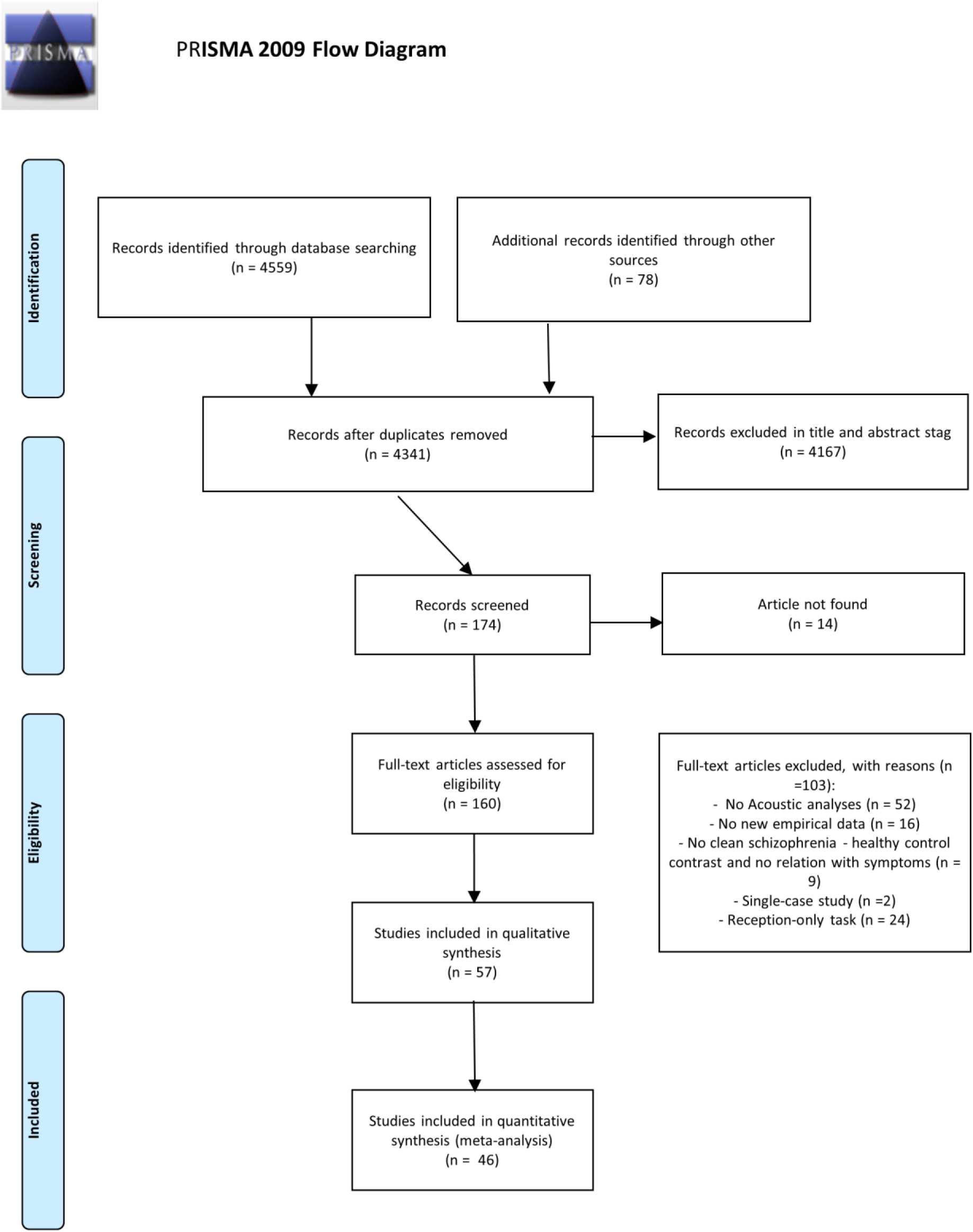
Flow chart showing the literature search and study selection process in accordance with the Preferred Reporting Items for Systematic Reviews and Meta-analyses (PRISMA) guidelines.

### Data extraction

For all the studies we reported the available clinical and demographic data, including pre-registered potential moderations. In particular we report: sample sizes, matching criteria, presence of a non-clinical control group, diagnosis, demographical variables (age, education, gender, language and ethnicity), clinical information (symptom clinical ratings, duration of illness, age of onset, hospitalization), level of intelligence (IQ), cognitive screening, medication. Further we extracted information about the speech production task, group-level acoustic estimates (mean and standard deviation), and correlation coefficients between acoustic measures and clinical ratings. While we extracted details about the speech production tasks (e.g. picture description, video description, biographical episodes), this fine-grained categorization involved too few studies in each category to be useful in a meta-analytic perspective. We therefore grouped speech production tasks into three broader categories: 1) *Constrained production* includes highly structured monological tasks such as reading aloud or repeating sequence of numbers. 2) *Free monological production* includes less constrained monological tasks such as description of pictures or videos, or providing narrative accounts (e.g. of a happy event, or of one’s life). Compared to constrained production, free production is more challenging, as the linguistic materials are less pre-defined by the task. 3) *Social interaction* includes structured and semi-structured interviews, as well as spontaneous conversations. The production is dialogical and involves interpersonal factors and dynamics. Compared to the other categories, social interaction is more challenging, as it both involves less pre-defined linguistic materials and the cognitive and social load of having to interact with another person. One could of course envision other categorization typologies; we therefore included all extracted task details in our publicly available dataset. Selected characteristics of included studies are available in Table 1.

**Table 1.**
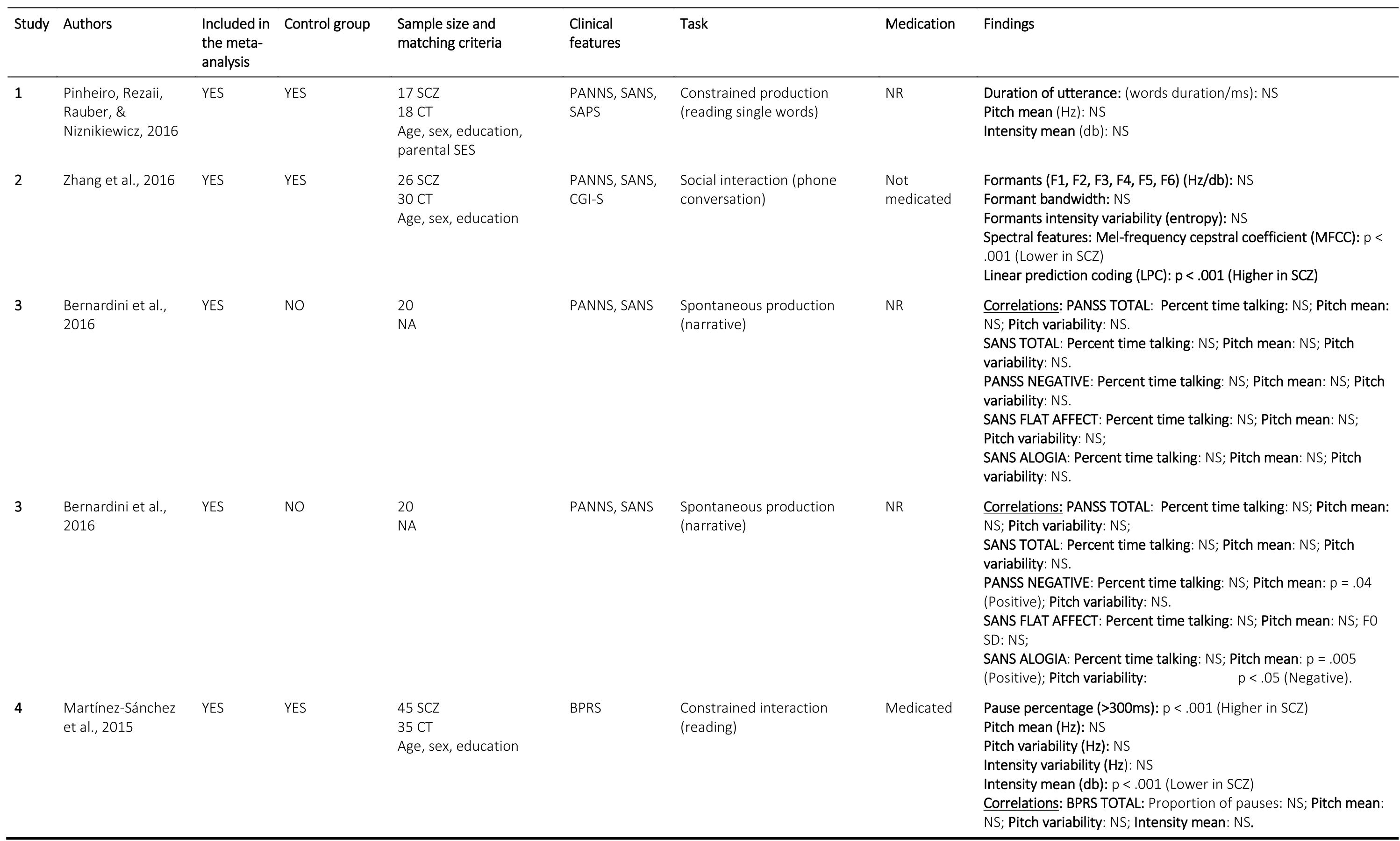
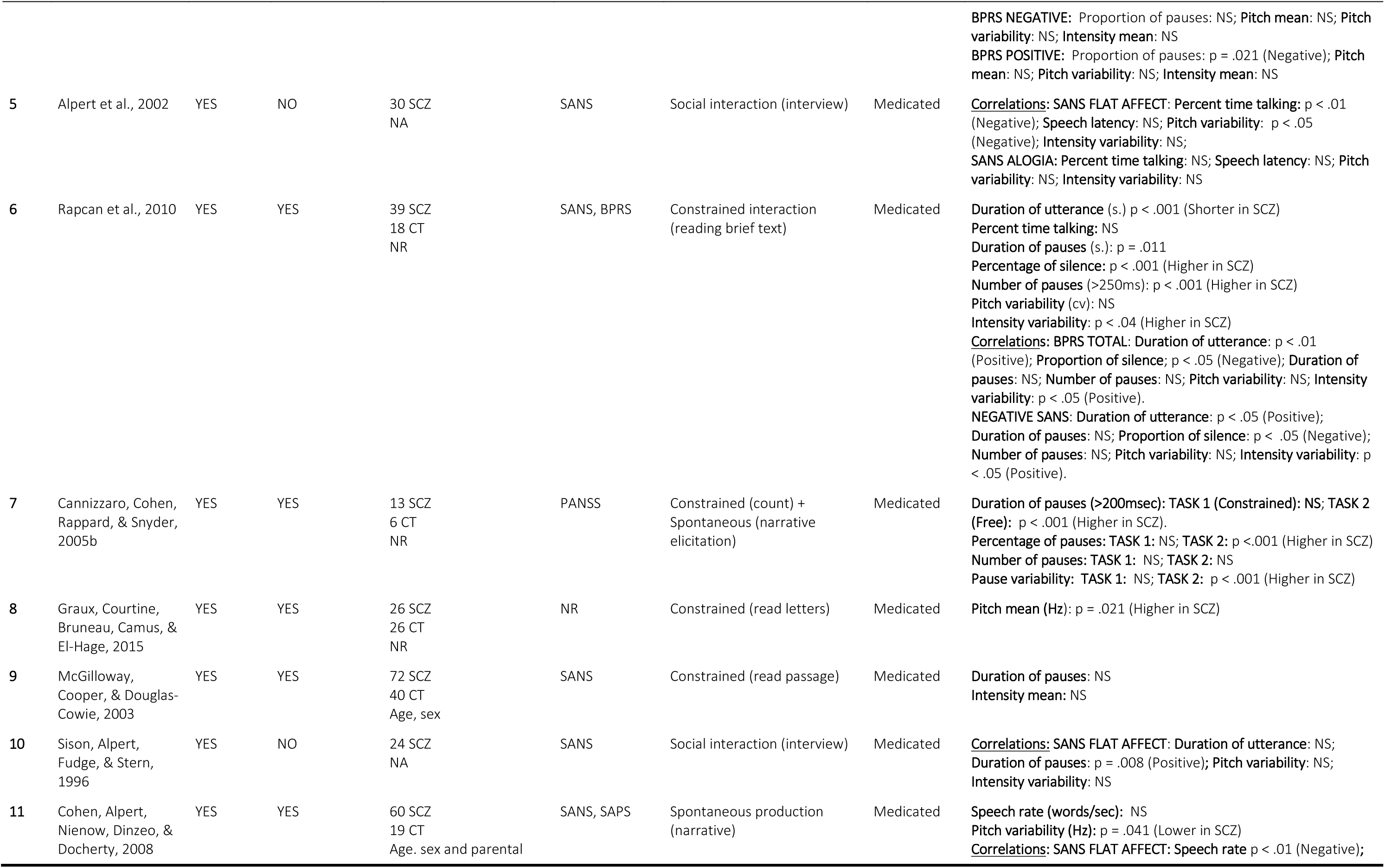
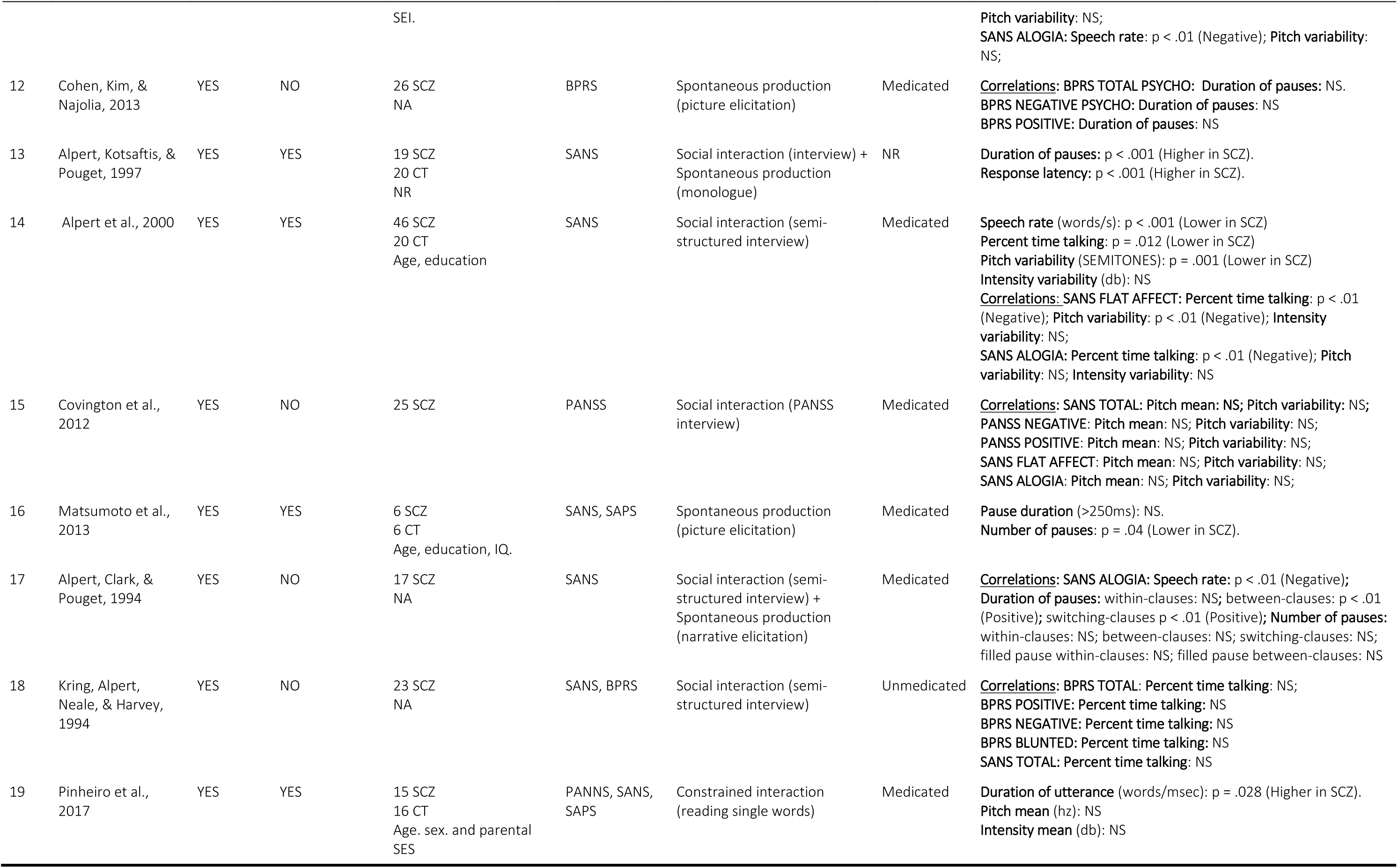
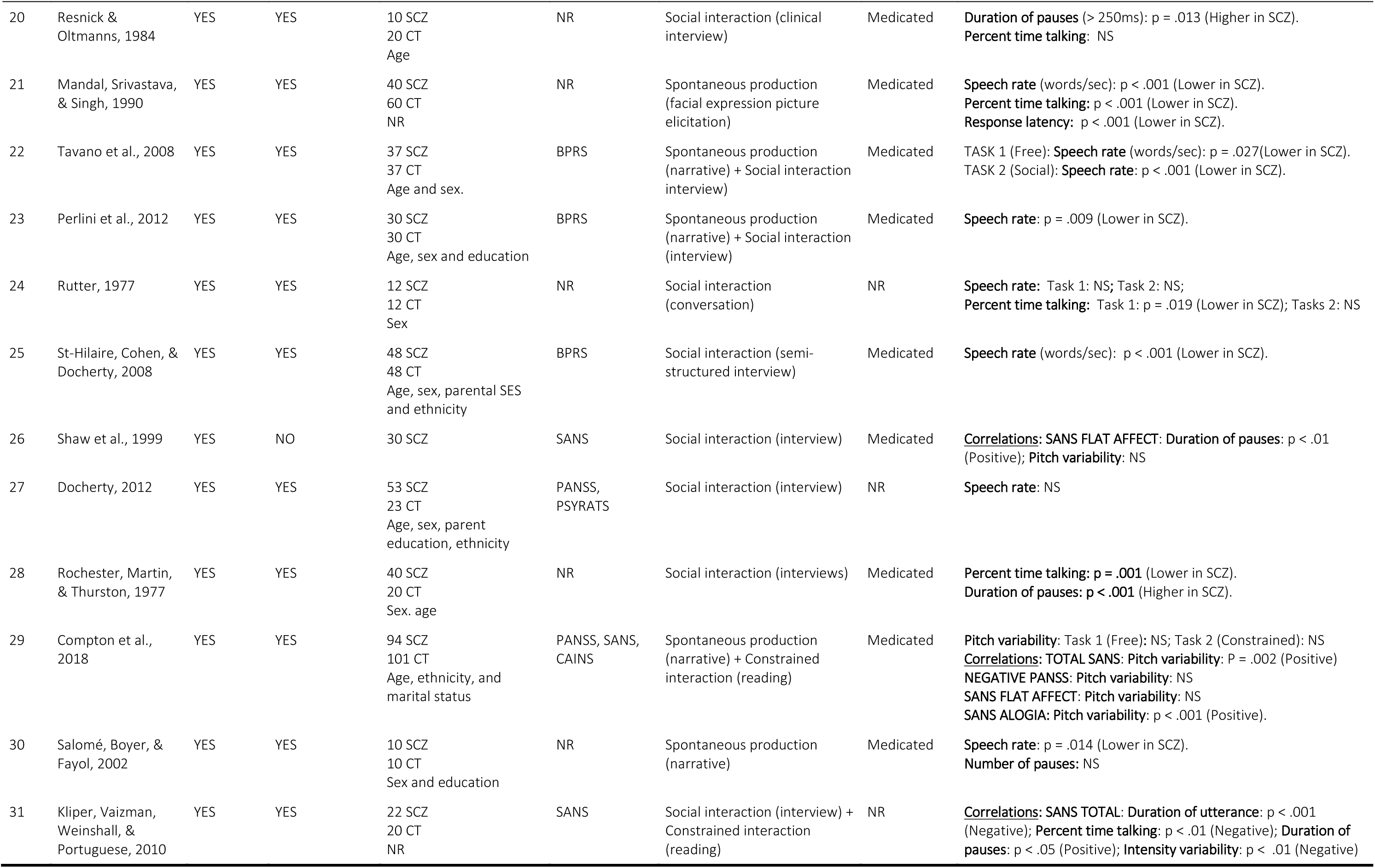
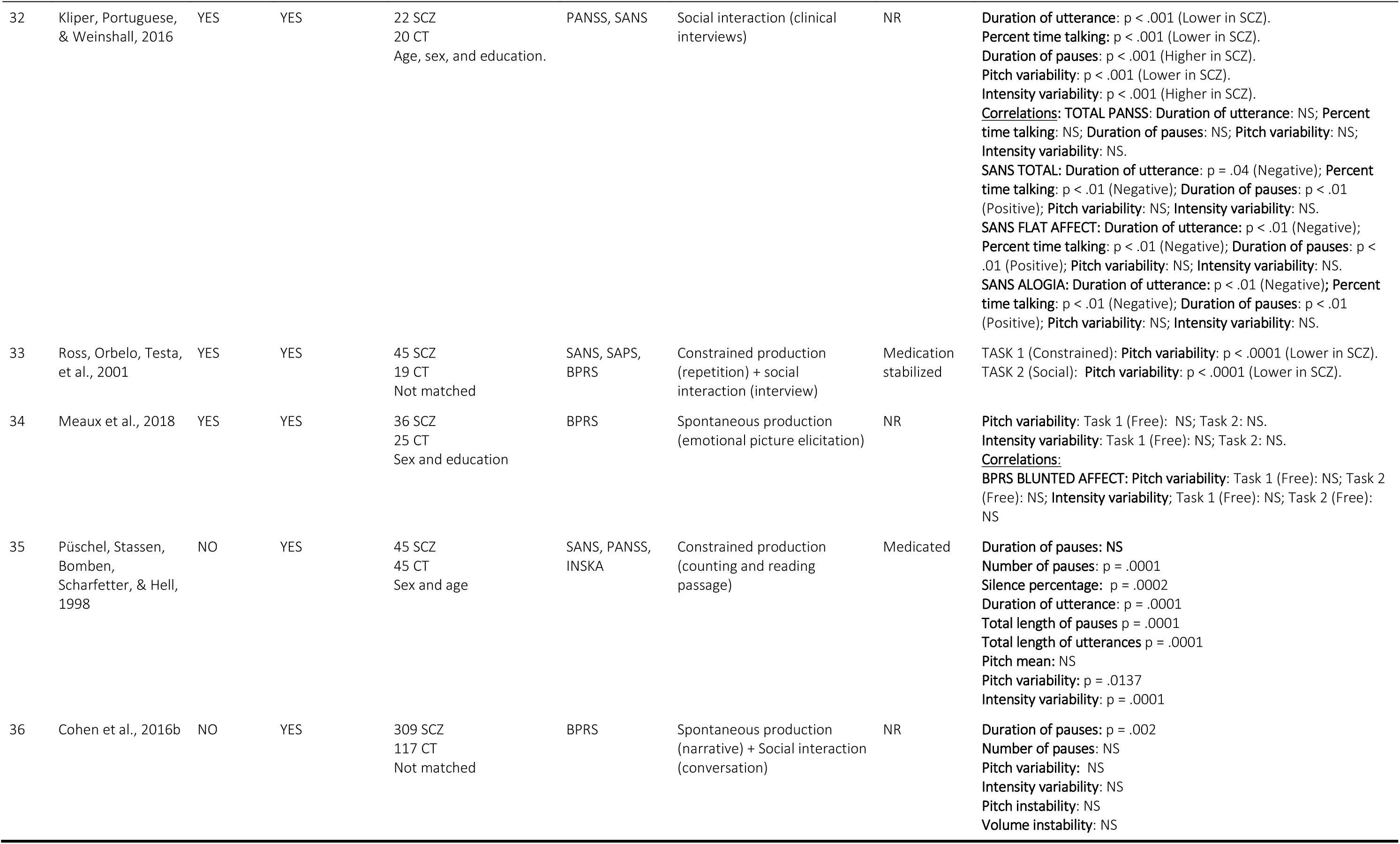
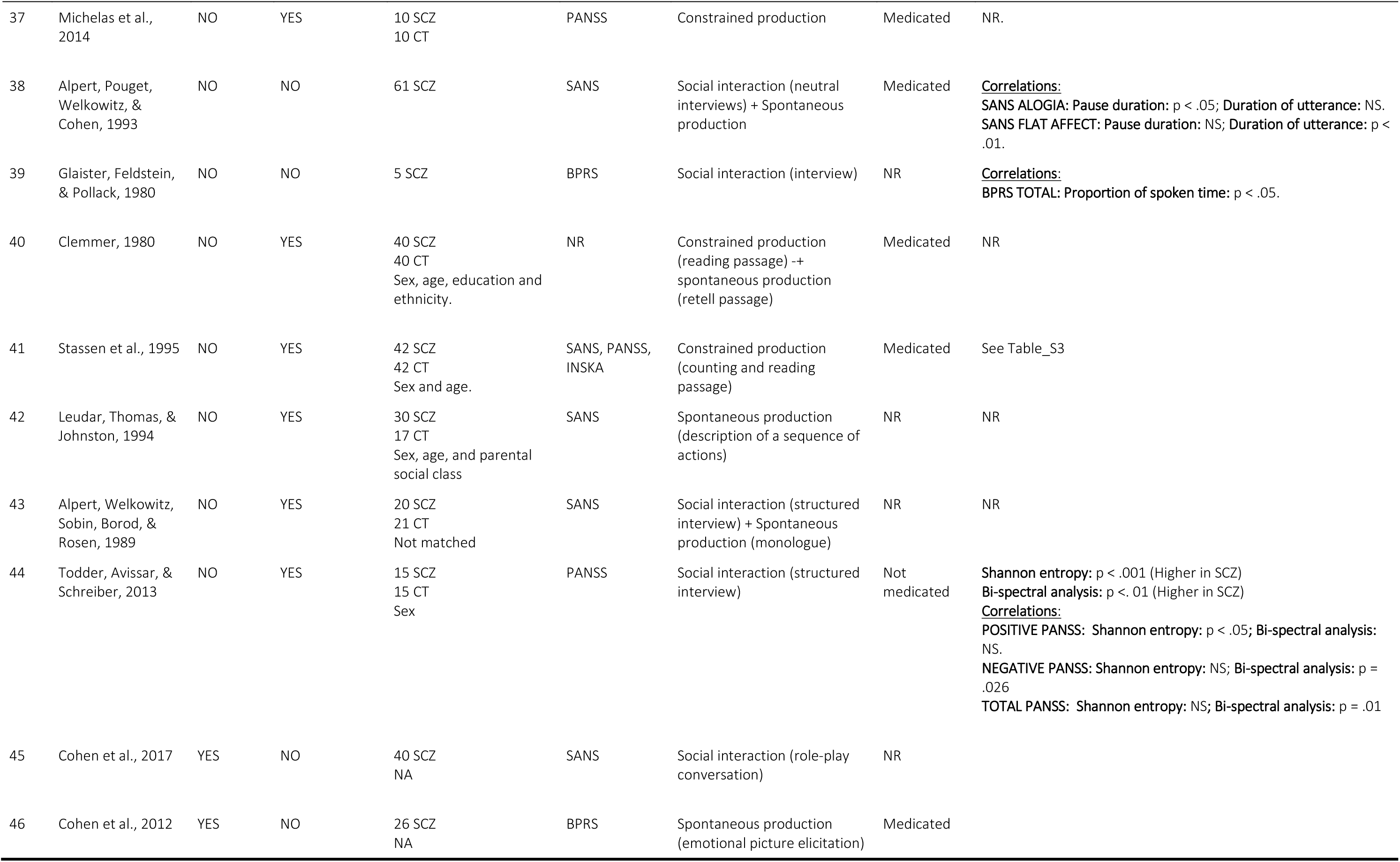
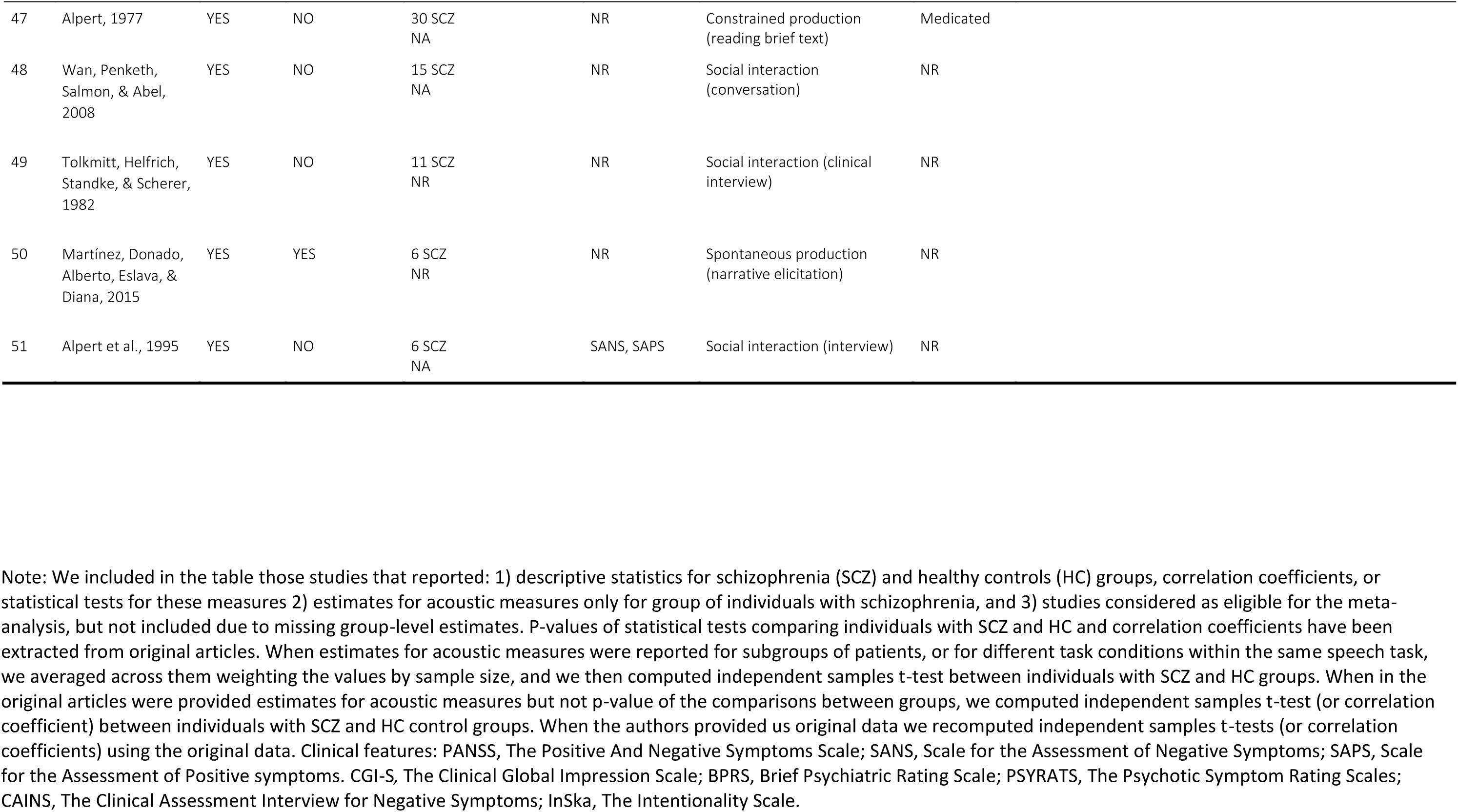
Selected characteristics of the studies included in the meta-analysis.

When more than four studies reported statistical estimates for an acoustic measure, or correlation with symptomatology, we performed meta-analysis of the effects. When estimates for acoustic measures were reported for subgroups of patients, or for different task conditions within the same speech task, we averaged across them weighting the values by sample size. In case of multivariate ML studies, we used a different focus. Multivariate ML approaches differ from the studies previously described in two main ways. While more traditional studies focus on a single feature at time, multivariate ML studies analyze multiple acoustic features simultaneously. While more traditional studies focus on best explaining all the current samples (minimizing within sample error), multivariate ML studies focus on generalizability of the results to new samples (minimizing out-of-sample error), e.g. by using validation and cross-validation techniques. In reviewing ML studies, we focused on reporting the algorithms adopted, the acoustic features considered and the performance of the algorithms in either discriminating individuals with schizophrenia from HC with respect to the acoustic measures considered or predicting the severity of clinical features (e.g. negative symptoms) from acoustic measures (see Table S3 in appendix).

We contacted all authors to obtain missing group-level estimates and individual-level data. Statistics on authors’ contact availability, propensity to respond and self-reported barriers to data sharing are also reported.

### Statistical analysis

Meta-analyses were performed following well-established procedures (Doebler and Holling, 2015; Field and Gillett, 2010; Quintana, 2015; Viechtbauer, 2010) and complemented by a Bayesian framework (Bürkner et al., 2017; Williams, Rast, & Bürkner, 2018). To estimate the differences in vocal patterns between individuals with schizophrenia and HC we extracted the standardized mean difference (SMD; also known as Hedges’ g). To estimate relations between vocal patterns and clinical features we extracted the raw correlation coefficient (Pearson’s r). These effects were analyzed using 2-level hierarchical Bayesian regression models to estimate the pooled effect sizes and corresponding credible (i.e., Bayesian confidence) intervals. The multilevel structure allowed us to explicitly model the heterogeneity (or σ^2^) in the results of the studies analyzed. By including a random effect by study, we assumed that the variability in experimental design, acoustic analyses and population samples could generate heterogeneous findings and allowed the model to estimate such heterogeneity. We then measured and tested for heterogeneity of the studies using the Cochran’s Q statistic (Cochran, 1954), which reveals how much of the overall variance can be attributed to true between-study variance. To analyze the influence of potential moderators explaining between study heterogeneity, meta-regression models were applied separately. Note that only speech task presented enough data points to be analyzed as moderator. Other pre-registered moderators were not sufficiently reported and would have required access to individual level data for adequate treatment.

Priors were chosen to be only weakly informative so that their influence on the meta-analytic estimates were small, only discounting extreme values: a normal distribution centered at 0 (no effect), with a standard deviation of 0.5 for the overall effect, and a positive truncated normal distribution centered at 0, with a standard deviation of 0.5 for the heterogeneity of effects (standard deviation of random effects). We report 95% credible intervals (CIs), i.e. the intervals within which there is a 95% probability that the true value of the parameter (e.g. effect size) is contained, given the assumptions of the model. We provide evidence ratios (ER) and credibility scores. ERs quantify the evidence provided by the posterior estimates in favor of the effect of diagnosis or of clinical feature (e.g. longer pauses in schizophrenia compared to HC) against the alternatives (e.g. same length or shorter pauses in schizophrenia). An ER equal to 3 indicates the hypothesis is 3 times more likely than the alternative. A credibility score indicates the percentage of posterior estimates falling above 0. Because Bayesian methods are less commonly used and understood, we also report p-values in order to reach a broader audience. Note that the p-values are calculated on the same 2-level hierarchical model as the Bayesian inference, with the difference that p-value statistics rely on completely flat priors and assume Gaussian distributions for all estimated parameters.

To assess the potential role of speech production task in explaining the patterns observed, we compared the baseline model with a second multilevel Bayesian model including task as predictor of difference in vocal patterns. We used Leave-One-Out-Information-Criterion (LOOIC) and stacking weights indicating the probability that the model including task is better able to predict new data than baseline (Vehtari et al., 2017).

To explore the possibility of publication bias, potential for funnel plot asymmetry was examined visually and tested using the rank correlation test (Begg and Mazumdar, 1994). The raw data and analysis scripts are available at https://osf.io/qdkt4/. The supplementary materials report an additional analysis including schizotypy. All computation was done in R(Core R Team, 2013) relying on metafor, brms and Stan (Burkner, 2013; Carpenter et al., 2017; Viechtbauer, 2010).

## Results

### 3.1 Study selection

See Fig. 1 for full details on the selection. We were able to retrieve relevant statistical estimates from 46 articles (55 studies) from the texts or the authors, with a total of 1254 patients (466 F) with schizophrenia and 699 controls (323 F). We contacted a total of 57 authors – including those of studies that were later deemed ineligible due to lack of statistical estimates – requesting additional information and individual level acoustic estimates for each participant: 40 (70.2%) responded and 10 (18%) provided at least some of the requested data. Chief reasons to decline sharing data were: i) effort required (n = 15, 50 %), ii) data loss (n = 14, 43.3% of respondents), iii) ethical concerns with data sharing (n = 3, 3 %), iv) skepticism towards quantitative meta-analyses (n =1, 3.3%). For more details on the email to the authors and their answers, see Supplementary Materials.

### 3.2 Differences in acoustic patterns between individuals with schizophrenia and healthy controls

Detailed results are reported in Table 2. Hierarchical Bayesian meta-analyses revealed significant effects of diagnosis (in terms of Hedges’ g) on pitch variability (-0.55, 95% CIs: - 1.06, 0.09), proportion of spoken time (-1.26, 95% CIs: -2.26, 0.25), speech rate (-0.75, 95% CIs: -1.51, 0.04), and duration of pauses (1.89, 95% CIs: 0.72, 3.21), see Fig. 2. No significant effect was found for pitch mean (0.25, 95% CIs: -0.72, 1.30), intensity variability (0.739, 95% CIs: -2.01, 3.39), duration of utterance (-0.155, 95% CIs: -2.56, 2.26) and number of pauses (0.05, 95% CIs: -1.23, 1.13). We generally found high heterogeneity between studies, indicating a likely high diversity in samples and methods, and publication bias, indicating a tendency to publish only significant results, thus making the published literature not fully representative of the actual population of study (see Table 2).

**Figure 2.**
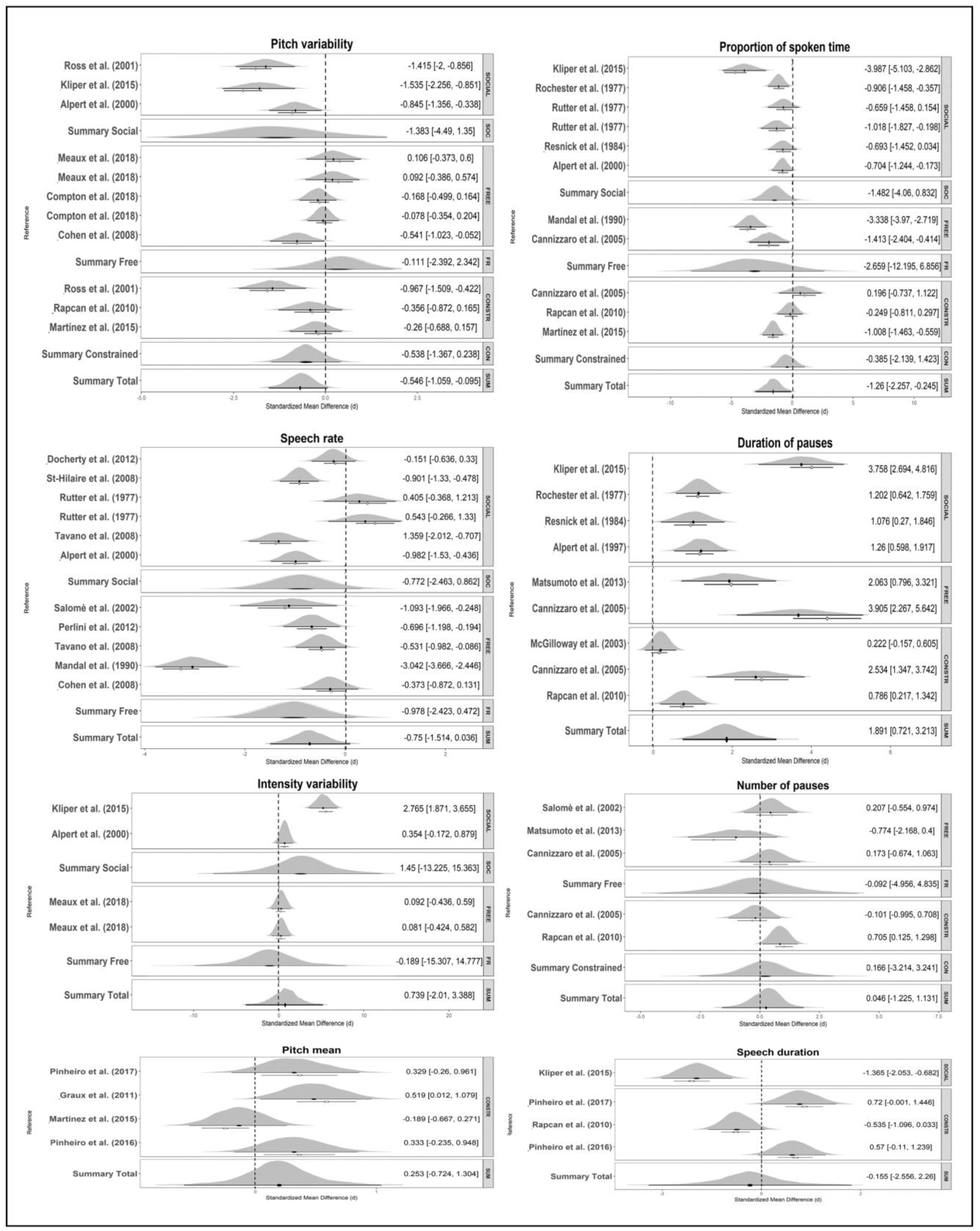
Forest plots of effect sizes (Hedges’g) for all the acoustic measures. The x-axis report effect sizes (black dot, positive values indicate that individuals with schizophrenia are higher on that acoustic measures, while negative values the opposite), posterior distribution (density plot) and original data point (white dot) for each study. The y-axis indicates the studies for which statistical estimates have been provided. The dotted vertical line indicates the null hypothesis (no difference between the populations). The studies are grouped by the speech task used to collect voice recordings (Constr = constrained monological, Free = free monological, Social = social interaction). When adding speech task credibly improved the model, we reported below each specific task group the summary effect size for that group. Filled diamonds represent summary effect sizes. Note that multiple inputs for the same reference may refer to original studies including: 1) more than one sample; 2) more than one speech production tasks (e.g. free monologue and dialogue); 3) more than one symptoms rating scale to measure symptomatology.

**Table 2.**
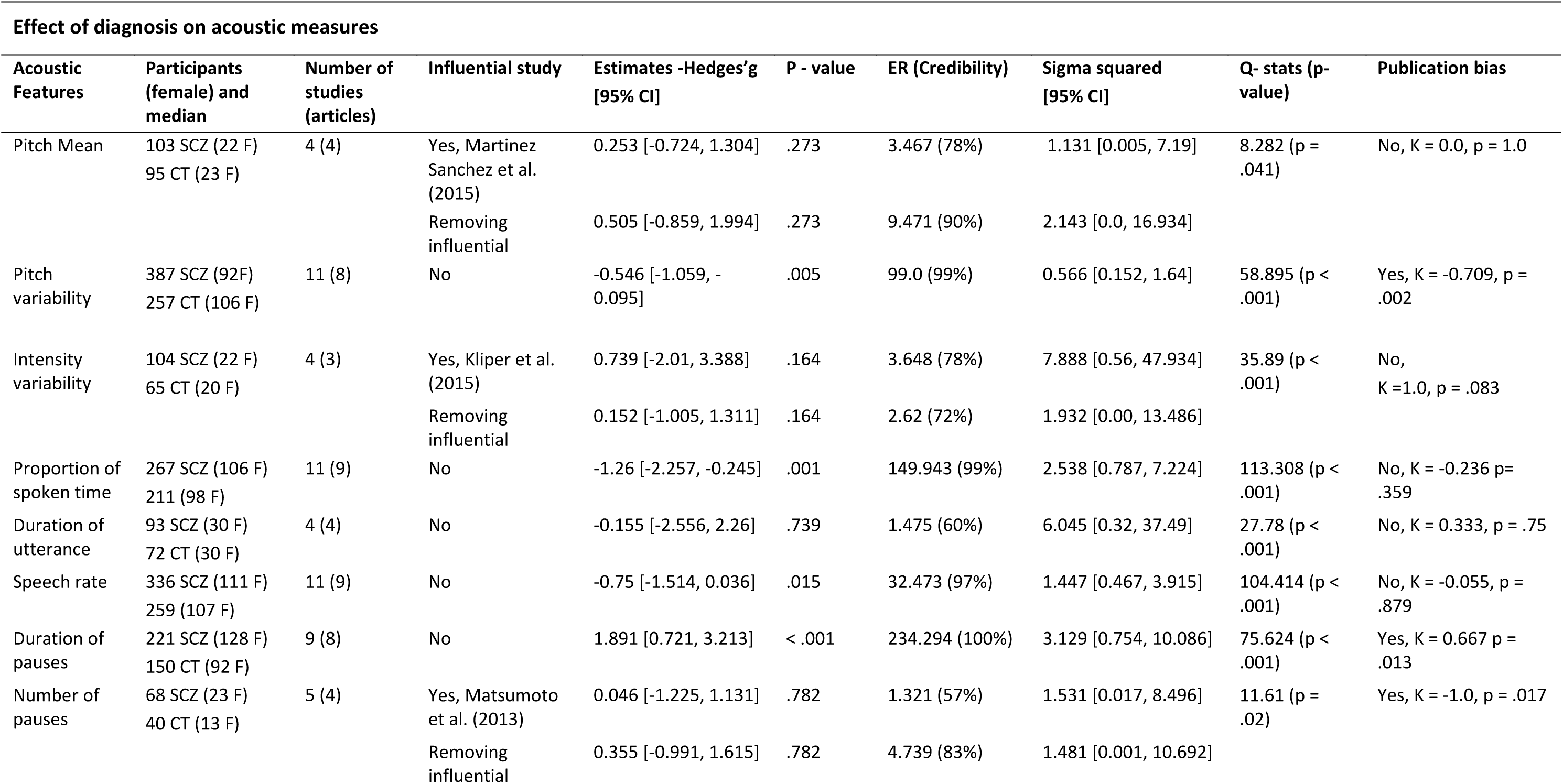
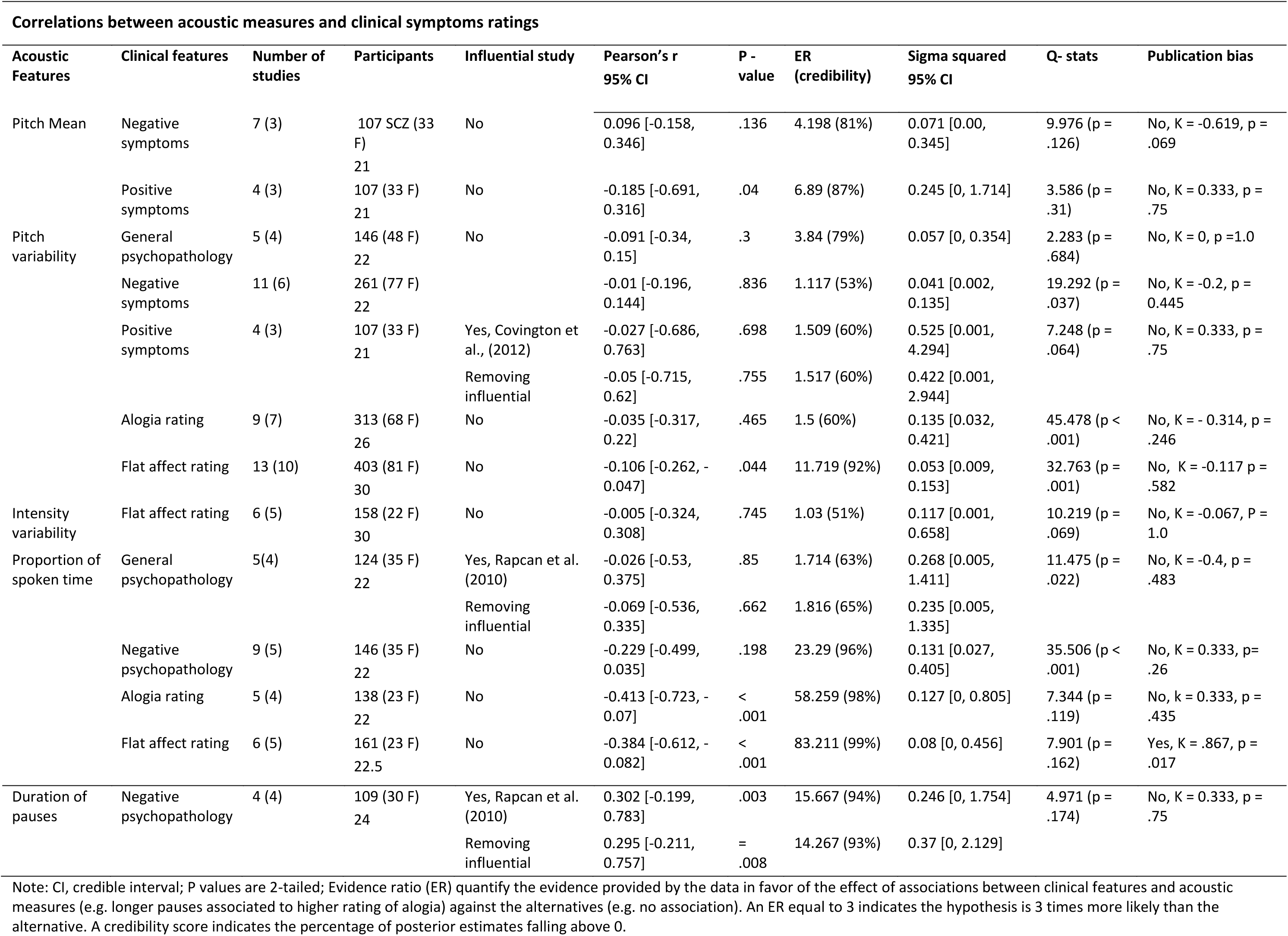
Main Results of the meta-analysis for the effect of diagnosis on acoustic measures, and for the correlations between acoustic measures and symptoms ratings.

| Effect of diagnosis on acoustic measures |  |  |  |  |  |  |  |  |  |
| --- | --- | --- | --- | --- | --- | --- | --- | --- | --- |
| Acoustic Features | Participants (female) and median | Number of studies (articles) | Influential study | Estimates -Hedges'g [95% CI] | P - value | ER (Credibility) | Sigma squared [95% CI] | Q- stats (p-value) | Publication bias |
| Pitch Mean | 103 SCZ (22 F)<br>95 CT (23 F) | 4 (4) | Yes, Martinez Sanchez et al. (2015) | 0.253 [-0.724, 1.304] | .273 | 3.467 (78%) | 1.131 [0.005, 7.19] | 8.282 (p = .041) | No, K = 0.0, p = 1.0 |
|  |  |  | Removing influential | 0.505 [-0.859, 1.994] | .273 | 9.471 (90%) | 2.143 [0.0, 16.934] |  |  |
| Pitch variability | 387 SCZ (92F)<br>257 CT (106 F) | 11 (8) | No | -0.546 [-1.059, -0.095] | .005 | 99.0 (99%) | 0.566 [0.152, 1.64] | 58.895 (p < .001) | Yes, K = -0.709, p = .002 |
| Intensity variability | 104 SCZ (22 F)<br>65 CT (20 F) | 4 (3) | Yes, Kliper et al. (2015) | 0.739 [-2.01, 3.388] | .164 | 3.648 (78%) | 7.888 [0.56, 47.934] | 35.89 (p < .001) | No, K = 1.0, p = .083 |
|  |  |  | Removing influential | 0.152 [-1.005, 1.311] | .164 | 2.62 (72%) | 1.932 [0.00, 13.486] |  |  |
| Proportion of spoken time | 267 SCZ (106 F)<br>211 (98 F) | 11 (9) | No | -1.26 [-2.257, -0.245] | .001 | 149.943 (99%) | 2.538 [0.787, 7.224] | 113.308 (p < .001) | No, K = -0.236 p = .359 |
| Duration of utterance | 93 SCZ (30 F)<br>72 CT (30 F) | 4 (4) | No | -0.155 [-2.556, 2.26] | .739 | 1.475 (60%) | 6.045 [0.32, 37.49] | 27.78 (p < .001) | No, K = 0.333, p = .75 |
| Speech rate | 336 SCZ (111 F)<br>259 (107 F) | 11 (9) | No | -0.75 [-1.514, 0.036] | .015 | 32.473 (97%) | 1.447 [0.467, 3.915] | 104.414 (p < .001) | No, K = -0.055, p = .879 |
| Duration of pauses | 221 SCZ (128 F)<br>150 CT (92 F) | 9 (8) | No | 1.891 [0.721, 3.213] | < .001 | 234.294 (100%) | 3.129 [0.754, 10.086] | 75.624 (p < .001) | Yes, K = 0.667 p = .013 |
| Number of pauses | 68 SCZ (23 F)<br>40 CT (13 F) | 5 (4) | Yes, Matsumoto et al. (2013) | 0.046 [-1.225, 1.131] | .782 | 1.321 (57%) | 1.531 [0.017, 8.496] | 11.61 (p = .02) | Yes, K = -1.0, p = .017 |
|  |  |  | Removing influential | 0.355 [-0.991, 1.615] | .782 | 4.739 (83%) | 1.481 [0.001, 10.692] |  |  |

VOICE IN SCHIZOPHRENIA: REVIEW AND META-ANALYSIS Correlations between acoustic measures and clinical symptoms ratings
| Acoustic Features | Clinical features | Number of studies | Participants | Influential study | Pearson's r<br>95% CI | P -<br>value | ER<br>(credibility) | Sigma squared<br>95% CI | Q- stats | Publication bias |
| --- | --- | --- | --- | --- | --- | --- | --- | --- | --- | --- |
| Pitch Mean | Negative symptoms | 7 (3) | 107 SCZ (33 F)<br>21 | No | 0.096 [-0.158, 0.346] | .136 | 4.198 (81%) | 0.071 [0.00, 0.345] | 9.976 (p = .126) | No, K = -0.619, p = .069 |
|  | Positive symptoms | 4 (3) | 107 (33 F)<br>21 | No | -0.185 [-0.691, 0.316] | .04 | 6.89 (87%) | 0.245 [0, 1.714] | 3.586 (p = .31) | No, K = 0.333, p = .75 |
| Pitch variability | General psychopathology | 5 (4) | 146 (48 F)<br>22 | No | -0.091 [-0.34, 0.15] | .3 | 3.84 (79%) | 0.057 [0, 0.354] | 2.283 (p = .684) | No, K = 0, p = 1.0 |
|  | Negative symptoms | 11 (6) | 261 (77 F)<br>22 |  | -0.01 [-0.196, 0.144] | .836 | 1.117 (53%) | 0.041 [0.002, 0.135] | 19.292 (p = .037) | No, K = -0.2, p = 0.445 |
|  | Positive symptoms | 4 (3) | 107 (33 F)<br>21 | Yes, Covington et al., (2012) | -0.027 [-0.686, 0.763] | .698 | 1.509 (60%) | 0.525 [0.001, 4.294] | 7.248 (p = .064) | No, K = 0.333, p = .75 |
|  |  |  |  | Removing influential | -0.05 [-0.715, 0.62] | .755 | 1.517 (60%) | 0.422 [0.001, 2.944] |  |  |
|  | Alogia rating | 9 (7) | 313 (68 F)<br>26 | No | -0.035 [-0.317, 0.22] | .465 | 1.5 (60%) | 0.135 [0.032, 0.421] | 45.478 (p < .001) | No, K = -0.314, p = .246 |
|  | Flat affect rating | 13 (10) | 403 (81 F)<br>30 | No | -0.106 [-0.262, -0.047] | .044 | 11.719 (92%) | 0.053 [0.009, 0.153] | 32.763 (p = .001) | No, K = -0.117 p = .582 |
| Intensity variability | Flat affect rating | 6 (5) | 158 (22 F)<br>30 | No | -0.005 [-0.324, 0.308] | .745 | 1.03 (51%) | 0.117 [0.001, 0.658] | 10.219 (p = .069) | No, K = -0.067, P = 1.0 |
| Proportion of spoken time | General psychopathology | 5(4) | 124 (35 F)<br>22 | Yes, Rapcan et al. (2010) | -0.026 [-0.53, 0.375] | .85 | 1.714 (63%) | 0.268 [0.005, 1.411] | 11.475 (p = .022) | No, K = -0.4, p = .483 |
|  |  |  |  | Removing influential | -0.069 [-0.536, 0.335] | .662 | 1.816 (65%) | 0.235 [0.005, 1.335] |  |  |
|  | Negative psychopathology | 9 (5) | 146 (35 F)<br>22 | No | -0.229 [-0.499, 0.035] | .198 | 23.29 (96%) | 0.131 [0.027, 0.405] | 35.506 (p < .001) | No, K = 0.333, p = .26 |
|  | Alogia rating | 5 (4) | 138 (23 F)<br>22 | No | -0.413 [-0.723, -0.07] | < .001 | 58.259 (98%) | 0.127 [0, 0.805] | 7.344 (p = .119) | No, k = 0.333, p = .435 |
|  | Flat affect rating | 6 (5) | 161 (23 F)<br>22.5 | No | -0.384 [-0.612, -0.082] | < .001 | 83.211 (99%) | 0.08 [0, 0.456] | 7.901 (p = .162) | Yes, K = .867, p = .017 |

|  |  |  |  |  |  |  |  |  |  |  |
| --- | --- | --- | --- | --- | --- | --- | --- | --- | --- | --- |
| Duration of<br>pauses | Negative<br>psychopathology | 4 (4) | 109 (30 F)<br>24 | Yes, Rapcan et al.<br>(2010) | 0.302 [-0.199,<br>0.783] | .003 | 15.667 (94%) | 0.246 [0, 1.754] | 4.971 (p =<br>.174) | No, K = 0.333, p =<br>.75 |
|  |  |  |  | Removing<br>influential | 0.295 [-0.211,<br>0.757] | =<br>.008 | 14.267 (93%) | 0.37 [0, 2.129] |  |  |

Note: CI, credible interval; P values are 2-tailed; Evidence ratio (ER) quantify the evidence provided by the data in favor of the effect of associations between clinical features and acoustic measures (e.g. longer pauses associated to higher rating of alogia) against the alternatives (e.g. no association). An ER equal to 3 indicates the hypothesis is 3 times more likely than the alternative. A credibility score indicates the percentage of posterior estimates falling above 0.

#### Moderator analysis

For detailed results, see Table S1 (in appendix). Adding the speech production task employed systematically increased the explained variability (stacking weights: 100%) in schizophrenia atypicalities for pitch variability (effects in Constrained Monologue, CM: -0.54; Free Monologue, FM: -0.11; Dialogue, D: -1.38), proportion of spoken time (CM: -0.38; FM: -2.7; D: -1.48), speech rate (CM: NA; FM: -0.98; D: -0.77), number of pauses (CM: 0.17; FM: - 0.09; D: NA), and intensity variability (CM: NA; FM: -0.19; D: 1.45). In general, we observe that dialogical and free speech show the biggest differences, while constrained monologue displays the smallest schizophrenia atypicalities in vocal patterns, except for pitch variability.

### 3.3 Correlation between acoustic measures and clinical ratings

For detailed results, see Table 2. Hierarchical Bayesian meta-analysis revealed significant overall correlation between flat affect and pitch variability (-0.11, 95% CIs: -0.26, 0.05) and proportion of spoken time (-0.38, 95% CIs: -0.61, -0.08), alogia and proportion of spoken time (-0.41, 95% CIs: -0.72, 0.07), positive symptoms and pitch mean (-0.19, 95% CIs: -0.69, 0.32), negative symptoms and pause duration (0.30, 95% CIs:-0.20, 0.78), see Fig. 3. No significant correlation was found between flat affect and intensity variability (-0.01, 95% CIs: -0.32, 0.31), alogia and pitch variability (-0.04, 95% CIs: -0.32, 0.22), general psychopathology and proportion of spoken time (-0.03, 95% CIs: -0.53, 0.375) and pitch variability (-0.09, 95% CIs:-0.34-, 0.15), positive symptoms and pitch variability (-0.03, 95% CIs: -0.68, .076), negative symptoms and pitch mean (0.01, 95% CIs:-0.16, 0.35), pitch variability (-0.01, 95% CIs:-0.20, 0.14), and proportion of spoken time (-0.23, 95% CIs: - 0.50, 0.04) (see Table 2). We generally found very high heterogeneity between studies, and publication bias, which suggests caution in interpreting these results and the need for more systematic replications (see Table 2).

**Figure 3.**
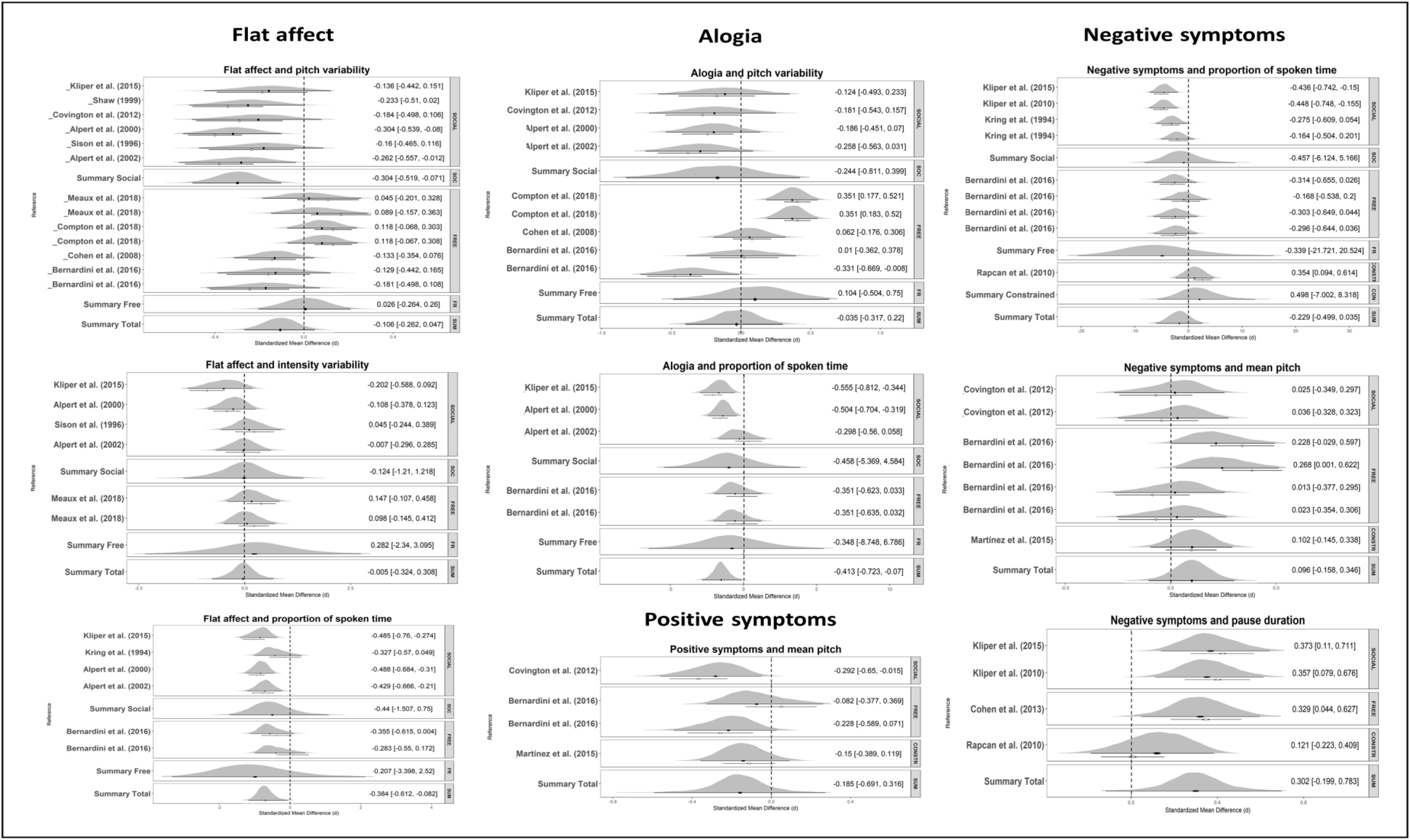
Forest plots of effect size (Pearson’s r) for the correlations between clinical symptoms and acoustic measures. The x-axis report effect sizes (black dot, positive values indicate a positive relation between acoustic measures and clinical symptoms rating, e.g. increased pause duration associated with increased rating of alogia, while negative values the opposite), posterior distribution (density plot) and original data point (white dot) for each study. The y-axis indicates the studies for which statistical estimates have been provided. The dotted vertical line indicates the null hypothesis (no difference between the populations). The studies are grouped as indicated in Figure 2

#### Moderator analysis

For detailed results, see Table S2 (in appendix). Adding speech task to the model credibly improved it (stacking weights: 100%) for correlations between *pitch variability* and positive symptom severity (effects in Constrained Monologue, CM: -0.001; Free Monologue, FM: - 0.40; Dialogue, D: 0.79), negative symptom severity (CM: 0.01; FM: 0.17; D: -0.43), alogia (FM: 0.10; D: -0.24) and flat affect (FM: 0.03; D: -0.30), between *intensity variability* and flat affect (FM: 0.28; D: -0.12), and between *proportion of spoken time* and total psychopathology (CM: 0.37; FM: -0.29; D: -0.15), negative symptom severity (CM: 0.49; FM: -0.34; D: -0.46), alogia (FM: -0.35; D: -0.46) and flat affect (FM: -0.21; D: -0.44). In general, we see that dialogic speech shows the strongest correlations with symptomatology, and constrained monological speech the weakest ones.

### 3.4. Multivariate machine learning (ML) studies

We found 5 ML articles fitting our criteria, all focused on identifying acoustic markers of the disorder (Kliper et al., 2016; Martínez-Sánchez et al., 2015; Püschel et al., 1998; Rapcan et al., 2010; Stassen et al., 1995) and 1 including the prediction of severity of clinical features from acoustic measures (Püschel et al., 1998). All acoustic features employed are reported in Table S3. Four studies employed linear discriminant analysis (LDA) and one employed support vector machines (SVM) to classify individuals with schizophrenia vs. HC. LDA linearly combines the acoustic features to generate a distribution of probability of a new voice belonging to an individual with schizophrenia. SVMs, on the contrary, construct multi-dimensional hyperplanes defined by the features and their interactions and there identify the regions that best separate the two groups. SVMs better account for non-linear relations between features and diagnosis. All studies reported accuracy beyond 75% and up to 87.5%. Results of four studies were cross-validated (Martínez-Sánchez et al., 2015; Rapcan et al., 2010; Stassen et al., 1995). Only one study (Rapcan et al., 2010) reported additional performance indices such as specificity, sensitivity, and area under the curve (AUC).

Two studies (Püschel et al., 1998; Stassen et al., 1995) attempted to predict the symptomatology (negative symptoms severity) from acoustic measures. The studies relied on LDA and reported an accuracy of 78.6% and 75.9% in classifying individuals with schizophrenia with higher vs lower scores of negative symptoms (PANSS negative < 11 or < 21 and SANS < 13 and < 37), and 71.4 % accuracy in predicting a future (14 days after) measurement of negative symptoms.

## Discussion

### Overview

Early descriptions of schizophrenia point to atypical voice patterns and studies relying on perceptual judgments and clinical ratings of voice patterns have indeed found large differences between patients and controls (Cohen et al., 2014). This suggests the existence of acoustic markers of the disorder. We set out to systematically review and meta-analyze the literature on the topic to assess the evidence for atypical acoustic patterns as markers of the disorder and to better inform future research. We were able to analyze the aggregated data from 46 unique articles including 1254 individuals with schizophrenia and 699 HC. The univariate studies identified several null results, as well as weak atypicalities in pitch variability (perhaps in relation to flat affect), and stronger atypicalities in duration (possibly related to alogia and flat affect). The effect sizes suggest a within-sample discriminative accuracy between 66% and 80%, likely less if assessing new data. The multivariate ML studies paint a more promising picture, with overall out-of-sample accuracies between 76.5% and 87.5%. When assessing the relation between acoustic features and symptomatology, we found that specific symptoms that are more directly related to voice, e.g. in their description in the clinical scales, yield slightly stronger results, with flat affect being related to speech variability and proportion of spoken time; and alogia being related to proportion of spoken time. Further, the results across all analyses suggest that dialogical productions, that is, tasks with a perhaps higher cognitive load and a more demanding social component, tend to involve larger effect sizes both in contrasting patients and controls and in assessing symptomatology. Free monological production follows and constrained production produces generally the smallest effects. Crucially, the studies analyzed mostly used widely different methods for sample selection, acoustic pre-preprocessing, feature extraction and selection. Indeed, we find large heterogeneity in the findings of the analyzed studies, and a large uncertainty in all our meta-analytic estimates.

#### What have we then learned?

In line with a previous non-systematic meta-analysis (13 studies, Cohen et al., 2014), we do indeed find evidence for acoustic markers of schizophrenia, further supporting the relation between clinical features of schizophrenia and voice patterns. However, the effect sizes are mostly too small for practical applications, not comparable to those of perceptual and clinical judgments, and in any case plagued by large between-studies variability. The only effect size which revealed to be large and robust was pause duration, specifically in relation with negative symptomatology, suggesting the potential of pause as a marker of schizophrenia symptomatology, which should be further investigated. While good progress has been made in the field, the review highlights a number of issues to be overcome to more satisfactorily understand acoustic patterns in schizophrenia and their potential. In particular we identified the following obstacles to the scientific understanding of acoustic features in schizophrenia: i) small sample sizes in terms of both participants and repeated measures, ii) heterogeneous, not fully up-to-date and underspecified methods in data collection and analysis, leading to scarce comparability between studies; iii) very limited attempts at theory driven research directly tackling the mechanisms underlying atypical vocal patterns in schizophrenia. These are discussed below.

#### Sample size

Schizophrenia is a heterogeneous disorder, and indeed several studies attempted to more specifically investigate the relation of acoustic features with the symptomatology of the disorder. However, given the limited meta-analytic effect sizes and the awareness that replications tend to show a marked shrinkage of effect sizes (Klein et al., 2018), we need to move beyond small heterogeneous studies. The majority of the studies analyzed include between 20 and 30 patients, plausibly due to the difficulty in accessing clinical populations. However, an expected Cohen’s d of 0.6 (pitch variability) would require at least 74 participants per group to reach a 95% power (calculations relying on G*Power (Faul et al., 2009)) at which effect size estimates are reliable (Schönbrodt and Perugini, 2013). If we considered the more conservative possibility of a smaller true effect size of 0.3, the required sample size would be 290 participants per group. While including as varied a sample as possible is an unavoidable concern, there are strategies to reduce the sample size needed. For instance, one could employ repeated measures, that is, collecting repeated voice samples over time. Using 10 repeated measures per participant brings the required sample from 290 participants per group to 82 (assuming that they are still representative of the full population). Repeated measures are also very useful to better understand the reliability of the acoustic patterns over re-testing and potentially across different contexts. In particular, we have seen that dialogical speech production tasks might yield stronger vocal differences, but without a controlled within-subject contrast it is difficult to assess whether this is due to the nature of the task or to other confounds in the sample and study design. We therefore recommend the use of multiple tasks systematically varying the cognitive and social demands involved in speech production.

Bigger sample sizes enable also a more fine-grained investigation of individual differences, such as the role of demographical (age, education, gender, language and ethnicity), cognitive and clinical features of the participants on their voice patterns and relatedly on the vocal markers of schizophrenia. Analyzing how acoustic features vary with fine-grained individual differences and context of speech production can help uncover the mechanisms behind atypical vocal patterns and provide an additional insight into schizophrenia. Indeed, we observe that acoustic features are more strongly related to specific symptoms (alogia, flat affect) than to global scores of psychopathologies.

#### Methods

We found that the field predominantly focuses on traditional acoustic features: pitch, intensity and duration measures. Even in these cases, the processing of the voice recordings and extraction of the features is poorly documented and arguably widely heterogeneous. Previous studies have found that different assumptions and settings in the feature extraction process might significantly affect the results (e.g. Kiss, van Santen, Prud’Hommeaux, & Black (2012) show different results for different choice of ceiling in pitch extraction). Further, speech pathology and speech signal processing research has developed a wide array of acoustic features more directly relatable to production mechanisms like fine-grained muscle control, or clarity of articulation (for some examples see Cummins, Sethu, Epps, Schnieder, & Krajewski, 2015), which are almost completely ignored in schizophrenia research. To overcome these barriers, we recommend the use of freely available open source software solutions providing standard procedures in the extraction of acoustic features and the documentation of the settings chosen (Boersma and Weenink, 2018; Degottex et al., 2014; Eyben et al., 2010)^2^. Use of new features should be compared against this baseline to facilitate comparability between studies.

Further, the vast majority of the studies focused on one acoustic feature at a time failing to produce effects comparable to those found in perceptual judgment studies. This supports the idea that perception is a complex process, non-linearly combining multiple acoustic cues. Multivariate techniques may thus allow to better capture vocal atypicalities. Indeed, the four ML studies we were able to identify provide promising out-of-sample accuracies, indicating that voice of individuals with schizophrenia may contain enough information to reliable distinguish between the two populations. However, the almost complete lack of overlap in features and methods employed in these studies makes it hard to assess how reliable the findings are across samples and whether there are more promising features and algorithms we should focus on.

#### Theory-driven research

A common feature of many of the studies reviewed is the lack of theoretical background. For example, limited attention is paid to clinical features and their severity and the choice of the speech-production task and acoustic measures used is often under-motivated. On the contrary, by putting hypothesized mechanisms to the test, more theory-driven research on vocal production in schizophrenia would improve our understanding of the disorder itself. For instance, social cognitive impairments (Green et al., 2015; Penn et al., 2008; Sergi et al., 2007) would motivate hypotheses on prosodic patterns when speaking to an interlocutor enabling the investigation of the impact of socio-cognitive mechanisms on vocal production, often hypothesized to be impaired in neuropsychiatric disorders. Comparison of narrative description and reading allows testing for the impact of word search (alogia) against a more general lack of motivation and energy (Frith, 1992; Lysaker and Bell, 1995; Trémeau et al., 2013), and adding a sustained phonation (say “aaaaa”) for the role of fine motor control of the vocal fold. Thus, using several tasks might help testing mechanistic hypotheses, and relatedly allow to assess whether speakers with a different clinical profile show differential vocal patterns across tasks. By including different tasks with diverse cognitive and social constraints, it would be possible to produce more robust results not specifically bound to a specific context, and to investigate the mechanisms and contextual factors responsible for voice abnormalities. Further, increased attention on the mechanisms should enable a more cross-diagnostic perspective, assessing the presence of atypicalities specific to schizophrenia or to more general cognitive and clinical features (Borsboom, 2017). For example, in other work we show that reduced pitch variability might be specific to schizophrenia, when compared to autism spectrum disorder and right hemisphere damage (Fusaroli et al., 2017; Weed and Fusaroli, 2019). However, we would expect to find reduced pitch variability also in affective disorders (Cummins et al., 2015b, 2015a, 2014). An approach based on clinical and cognitive profiles would provide more informative support to the clinicians than just the identification of the diagnostic class.

Further, little work has been done on the physiological mechanisms underlying atypical voice production in schizophrenia, e.g. whether related to auditory processing, pitch control, neuromotor disorders, or antipsychotic medications (Cannizzaro et al., 2005; Konopka and Roberts, 2016; Matsumoto et al., 2013; Peluso et al., 2012; Tenback et al., 2010; Walther, 2015; Walther and Strik, 2012). This is an important venue for further investigations.

Finally, speech involves not only voice but also more linguistic aspects: for instance, lexical choices, syntactic and semantic structure are all mentioned in the symptomatology of the disorder (e.g. poverty of content and tangentiality). Investigating these additional features of speech complementing acoustic analyses could yield further insight. For example, recent research (Çokal et al., 2019) showed that only syntactically motivated and not general pause patterns reliably distinguished participants with schizophrenia from non-clinical controls. Further, automated measures of semantic coherence are being developed to assess symptoms like tangentiality (Corcoran et al., 2018). It is an open question how they relate to vocal patterns and how they can complement each other.

#### Open Science

The recommendations to rely on large sample sizes, include individual differences, and cumulatively employ acoustic features from previous studies might seem too cumbersome, or even unreasonable, given the high costs of research, ethical and practical constraints in accessing clinical populations and proliferation of acoustic measures. This is why we recommend open science practices to be included already in the research design. Releasing in controlled and ethically sound ways one’s datasets enables the construction of large collective samples and re-analysis of the data to replicate and extend previous findings. Accessing previous datasets is currently unfeasible, due to lack of answers from corresponding authors, data loss and the practical and time-consuming hurdle of finding, preparing and sharing the data years after the study has been published. This suggests that planning data-sharing from the onset of the study is necessary to ensure a more open, collective and nuanced science of acoustic markers in schizophrenia, conscious of the individual differences and diverse symptomatology. Sharing identifiable (voice) data related to clinical populations requires serious ethical considerations and careful sharing systems, but there are available datasets of voice recordings in e.g. people with Parkinson’s, bipolar disorder, depression and autism spectrum disorder (Ambite et al., 2015; Cummins et al., 2015b; Gratch et al., 2014; Schuller et al., 2013; Tsanas et al., 2014), thus suggesting that these hurdles can be overcome. In line with these recommendations, all the data and the codes used in this manuscript are available at https://osf.io/qdkt4/.

## Conclusion

We have systematically reviewed the evidence for acoustic markers of schizophrenia and its symptomatology, as well as the research practices employed. We did not find conclusive evidence for clear acoustic markers of schizophrenia, although pitch variability and duration are potential candidates, with strong but heterogeneous evidence in favor of the use of pause duration. Multivariate studies are more promising, but their generalizability across samples could not be assessed. To advance the study of vocal markers of schizophrenia we outlined a series of recommendations towards more cumulative, open, and theory-driven research.

## Supporting information

SupplementaryMaterials_Schizotypy

SupplementaryMaterials_Mail_analysis

SupplementaryMaterial_RiskOfBias

Table_S1_Moderator_analysis_Contrast_SM

Table_S2_Moderator_analysis_Correlations_SM

Table_S4_Feature_description_SM_060819_RESUB

Table_S5_Results_meta_analysis_schizotypy_SM

Table_S3_Machine_Learning_SM

## Footnotes

1 We included schizotypy in literature search to better cover schizophrenia spectrum disorder. However, given schizotypy is only included in the schizophrenia spectrum in the ICD and is mentioned in the personality disorders in the DSM classification, we only included schizotypy in additional analysis in the supplementary material. Studies without a control group, i.e. only including patients with schizophrenia, were included in the qualitative synthesis and data extraction as possible additional datapoints for future studies.

2 Note that while the bibliographic references date to 2010 and 2014, the referred software has been frequently updated since.

## Notes

#### Summary of Updates

Revisions to the manuscript and SM.

https://osf.io/qdkt4/

