## SupplementaryMaterials_Schizotypy for "Voice Patterns in Schizophrenia: A systematic Review and Bayesian Meta-Analysis"

**Supplementary analysis including studies of schizotypy**

**Introduction**

Schizotypy is defined as the personality organization reflecting a latent liability for schizophrenia with an incidence of approximately 10%^1–3^. The schizophrenic phenotype can result in various psychological and behavioral manifestations, such as subtle thought disorder or excessive interpersonal fear. Schizotypy is a useful theoretical construct that offers a broad continuum along which different clinical and subclinical manifestations can be unified for understanding the psychopatology associated with the schizophrenia spectrum^4^. To provide a more complete picture of the acoustic markers of schizophrenia, we here provide an additional analysis including the studies of acoustic markers in schizotypy^[1]^ and assessing whether this personality organization present analogous patterns to schizophrenia.

**Methods**

We used the same methods as in the main manuscript. As in the analysis of moderators, we assessed whether distinguishing between schizophrenia and schizotypy improved our ability to explain the data, using a LOOIC based model comparison against the null model not discriminating between the two.

**Results**

The meta-analysis included 55 studies (46 articles), of which 6 (5 articles) involved participants with schizotypy for a total of 1254 participants with schizophrenia (466 female ones), 202 participants with schizotypy (136 female ones) and 859 comparison participants (428 female ones). Detailed results are reported in Table 7.

When analyzed in isolation, individuals with ST displayed credible and significant differences from HC in number of pauses only (1.398, 95% CIs: -1.086, 3.778). However, hierarchical Bayesian meta-analyses revealed that schizophrenia and schizotypy present credible and significant differences in their vocal atypicalities for what regards pitch and intensity variability, speech rate and duration of pauses. In other words, while schizophrenia involves a lower pitch variability than HC, schizotypy involves credibly higher pitch variability (difference in effects between SZ and ST: -1.577, 95% CIs: -3.076, -0.069). Where schizophrenia involves more pronounced swings in intensity than HC, schizotypy displays relatively less pronounced swings (1.872, 95% Cis: -2.109, 5.916). Where schizophrenia involves slower speech rate and longer pauses than HC, schizotypy presents credibly faster speech rate and shorter pauses (respective differences: -2.364, 95% CIs -5.048, 0.266; and 3.402, 95% CIs: -0.392, 7.276). No credible and significant differences were found for pitch mean, proportion of spoken time, duration of utterance, number of pauses.

Heterogeneity between studies was high and could not be reduced to random sample variability for of pitch mean, pitch variability, mean intensity, intensity variability, proportion of spoken time, speech rate, duration of utterance, number of pauses and duration of pauses, indicating a likely high diversity in samples, and methods.

Publication bias was found for duration of pauses, indicating for these measures a tendency to publish only results that show a significant finding, thus making the published literature not fully representative of the actual population of studies.

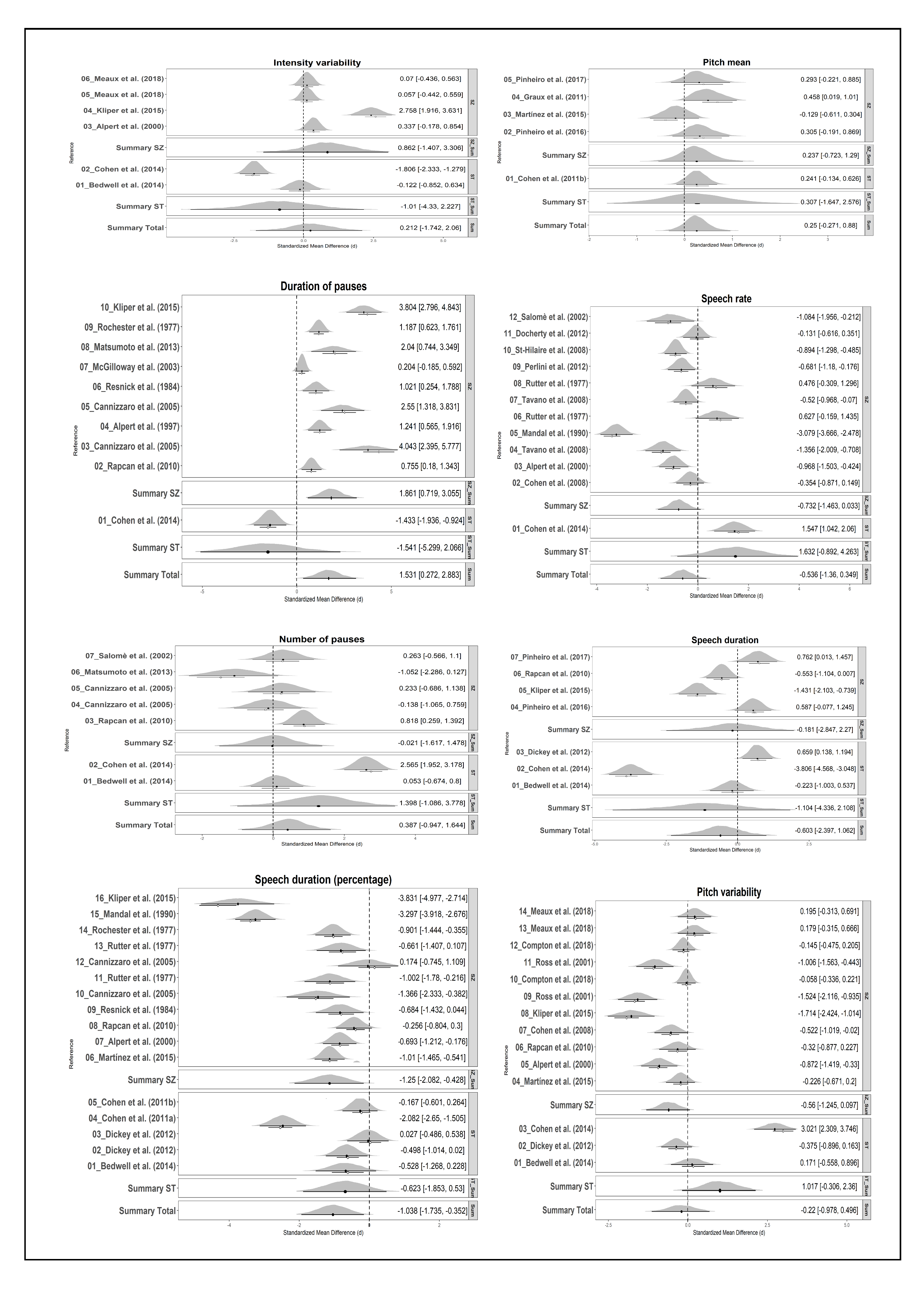

Fig. 7 Forest plots of effect sizes (Hedges’g) for all the acoustic measures meta-analyzed. The x-axis report effect sizes (black dot, positive values indicate that individuals with SCZ or ST are higher on that acoustic measures, while negative values the opposite), posterior distribution (density plot) and original data point (white dot) for each study. The y-axis indicates the studies for which statistical estimates has been provided. The dotted vertical line indicates the null hypothesis (no difference between the populations). The studies are grouped by the diagnosis (SCZ or ST). Filled diamonds represent summary effect sizes.

**Discussion**

We found that while evidence for atypical vocal patterns (in relation to controls) in schizotypy is limited, with only credible and significant differences in the number of pauses, more robust differences are observed when comparing atypicalities in schizotypy to those in schizophrenia. These differences tend to be in opposite directions: where schizophrenia has slower speech, lower pitch swings and higher intensity swings than controls (see also Table 2 in main manuscript), schizotypy displays differences in the opposite patterns. These findings do not support the idea of a continuum between SZ and ST in vocal abnormalities, with vocal features of ST more similar to HC than to SZ.

The only possible exception to this is the proportion of spoken time, for which schizophrenia displays markedly lower proportion than HC (-1.25, 95% Cis: -2.082 -0.428), while schizotypy a more moderate but still negative difference in proportion (0.623, 95% Cis -1.853, 0.53), thus potentially indicating a continuum.

An important limitation to these analyses of schizotypy is the very limited number of studies available: 5 studies for proportion of spoken time, 3 studies for pitch variability, 2 for intensity variability, and only 1 for speech rate and pause duration involving participants with schizotypy. In particular, credible and significant differences were biased by a single study (Cohen et al. 2014) for all the four measures considered.

Our results thus do not support the hypothesis of a general continuum. However, considering the large heterogeneity we found in all the findings, and the large uncertainty in all our meta-analytic estimates, the evidence available is largely insufficient to a rigorous test of the hypothesis and more work is needed.
