## SupplementaryMaterials_Mail_analysis for "Voice Patterns in Schizophrenia: A systematic Review and Bayesian Meta-Analysis"

**Supplementary analysis on data availability**

**Table 8. Data Availability by Year of Publication**

| **PERIOD** | **MAIL_SENT** | **NOT_FOUND_NO WORKING E-MAIL** | **RESPONSE TO E-MAIL** | **NO RESPONSE TO E-MAIL** | **DATA_PROVIDED** | **RESPONSE BUT DATA NOT PROVIDED** | **MOTIVATIONS FOR NOT PROVIDING RESEARCH DATA, GIVEN BY THE ORIGINAL AUTHORS:** | | | | **DATA EXTANT** | **NUMBER OF PAPERS** |
| --- | --- | --- | --- | --- | --- | --- | --- | --- | --- | --- | --- | --- |
|  |  |  |  |  |  |  | **DATA LOST OR NOT COLLECTED** | **TOO MUCH EFFORT REQUIRED** | **ETHICAL_CONCERNS** | **SKEPTICISMS** |  |  |
| 1960-1965 | 0 | 0 | 0 | 0 | 0 | 0 | 0 | 0 | 0 | 0 | 0 | 0 |
| 1966-1970 | 0 | 2 (100%) | 0 | 0 | 0 | 0 | 0 | 0 | 0 | 0 | 0 | 2 |
| 1971-1975 | 0 | 0 | 0 | 0 | 0 | 0 | 0 | 0 | 0 | 0 | 0 | 0 |
| 1976-1980 | 1 | 4 (80%) | 1 (100%) | 0 | 0 | 1 (100%) | 1 (100%) | 0 | 0 | 0 | 0 (0%) | 5 |
| 1981-1985 | 2 | 2 (50%) | 1 (50%) | 1 (50%) | 0 | 1 (100%) | 1 (100%) | 0 | 0 | 0 | 0 (0%) | 4 |
| 1986-1990 | 2 | 0 (0%) | 2 (100%) | 0 | 0 | 2 (100%) | 2 (100%) | 0 | 0 | 0 | 0 (0%) | 2 |
| 1991-1995 | 6 | 1 (14.2%) | 6 (100%) | 0 | 0 | 6 (100%) | 4 (66.6%) | 1 (16.6%) | 0 | 1 (16.6%) | 2 (33.3%) | 7 |
| 1996-2000 | 3 | 2 (40%) | 3 (100%) | 0 | 0 | 3 (100%) | 2 866.6%) | 1 (33.3%) | 0 | 0 | 1 (33.3%) | 5 |
| 2001-2005 | 7 | 0 | 5 (71.4%) | 2 (28.6%) | 2 (40%) | 3 (60%) | 3 (100%) | 0 | 0 | 0 | 2 (28.5%) | 7 |
| 2006-2010 | 10 | 0 | 4 (40%) | 6 (60%) | 0 | 4 (100%) | 1 (25%) | 3 (75%) | 0 | 0 | 3 (30%) | 10 |
| 2011-2015 | 19 | 0 | 12 (63.1%) | 7 (36.9%) | 4 (33.3%) | 8 (66.6%) | 1 (12.5%) | 6 (75%) | 1 (12.5%) | 0 | 11 (57.9%) | 19 |
| 2016-2020 | 7 | 0 | 6 (85.7%) | 1 (14.3%) | 4 (66.6 %) | 2 (33.3%) | 0 | 2 (100%) | 0 | 0 | 6 (85.7%) | 7 |
| **Total** | 57 | 11 (16.2%) | 40 (70.2 % | 17 (29.8%) | 10 (25%) | 30 (75%) | 15 (50%) | 13 (43.3%) | 1 (3.3%) | 1 (3.3%) | 25 (43.9%) | 68 |

**Table legend**

PERIOD: Age period

MAIL_SENT: Number of e-mail sent to original authors

NOT_FOUND_NO WORKING E-MAIL: Number of papers for which we were not able to find a working e-mail. Percentage is calculated as: NOT_FOUND_NO WORKING E-MAIL/(MAIL SENT+ NOT_FOUND_NO WORKING E-MAIL)

RESPONSE TO E-MAIL: Number of papers for which we received an e-mail response from the original authors. Percentage is calculated as: RESPONSE TO E-MAIL/MAIL_SENT

NO RESPONSE TO E-MAIL: Number of papers for which we have not received an e-mail response from the original authors. Percentage is calculated as: NO RESPONSE TO E-MAIL/MAIL_SENT

DATA_PROVIDED: Number of papers for which we have received at least some of the requested data from the original authors. Percentage is calculated as: DATA_PROVIDED/RESPONSE TO E-MAIL

RESPONSE BUT DATA NOT PROVIDED: Number of papers for which we have received an e-mail response from the original authors, but the authors have not provided us at least some of the requested data. Percentage is calculated as: RESPONSE BUT DATA NOT PROVIDED /RESPONSE TO E-MAIL

**Motivations for not providing research data, given by the original authors:**

1) DATA LOST OR NOT COLLECTED: Number of papers for which data were lost, not found or no longer accessible by the original authors. Percentage is calculated as DATA LOST OR NOT COLLECTED/DATA_PROVIDED
2) TOO MUCH EFFORT REQUIRED: Number of papers for which data exists, but it requires too much effort to retrieve the data. Percentage is calculated as TOO MUCH EFFORT REQUIRED /DATA_PROVIDED
3) ETHICAL_CONCERNS: Number of papers for which data sharing is not possible due to ethical constraints. Percentage is calculated as ETHICAL_CONCERNS /DATA_PROVIDED
4) SKEPTICISMS: Number of papers for which authors did not agree to share data due to skepticisms toward our meta-analylsis

DATA EXTANT: Number of papers for which data are extant. Percentage is calculated as: (TOO MUCH EFFORT REQUIRED+ ETHICAL_CONCERNS+ SKEPTICISMS + DATA_PROVIDED)/MAIL SENT

NUMBER OF PAPERS: Total number of papers. The total number of papers exceeds the number of papers considered eligible for the meta-analysis, because some of the papers that we considered eligible in a first stage, or for which we were in doubt about eligibility (and for which we have thus sent e-mail to original authors), were in a second stage no longer considered as eligible. We included these papers and authors in this analysis, because they are totally representative of the entire population of the researchers within this research field.

**Statistical analysis on data availability**

We examined the effect of years since article was published on obstacles to receiving data from the authors (for a similar analysis see Vines et al., 2014^1^). We used logistic regression to formally investigate the relationships between the age of the paper and: 1) probability to find a working email 2) probability of receiving response from authors given that at least one email was working 3) probability that original data were still extant (i.e. authors provided us the data, or answered that they were not able to provide us the data due to reasons different from data not extant) 4) probability to receive at least some of the requested data given that authors replied to the mail sent.

1) Probability to find at least one working working email

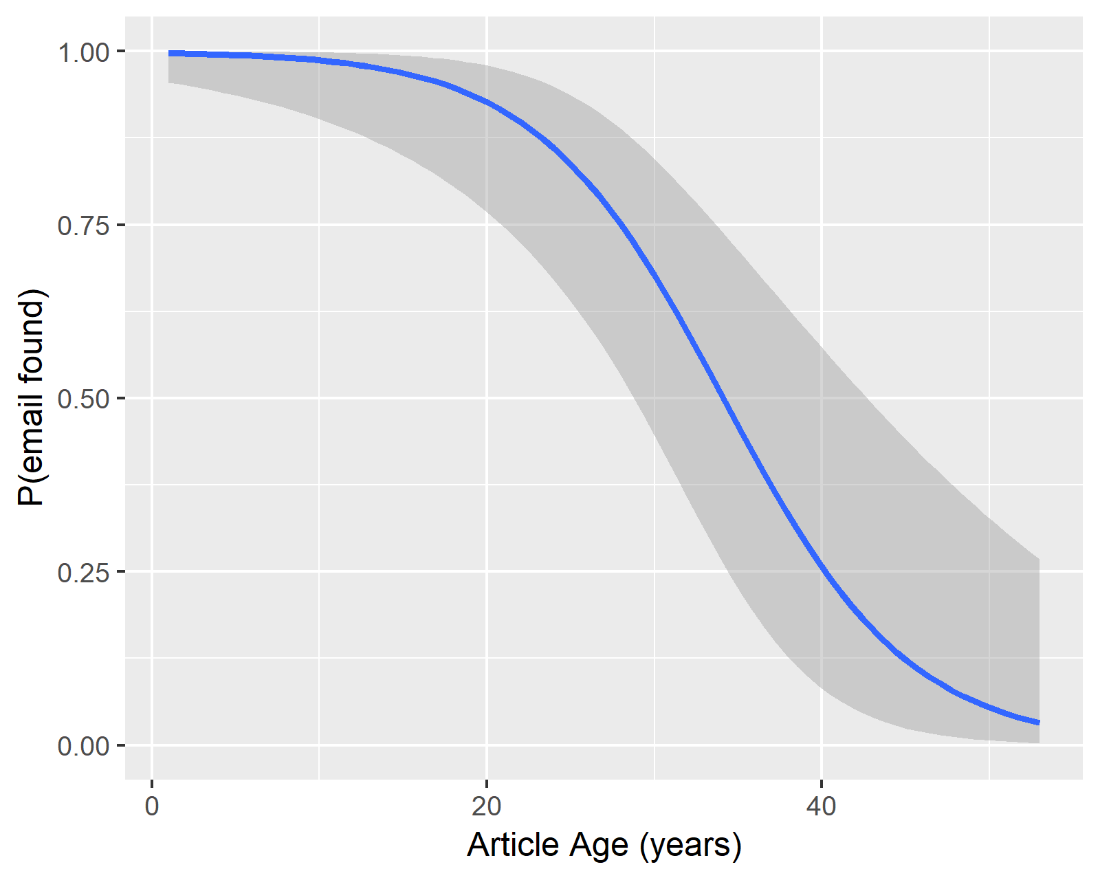

Logistic regression coefficients (se):

Intercept	6.135 (1.526), z-value = 4.020, p < .001.

Article age (years)	-0.180 (0.048), z-value = -3.714, p <. 001.

The probability to found at least one working e-mail declines rapidly with article age.

2) Probability of receiving response from authors given that at least one email was working

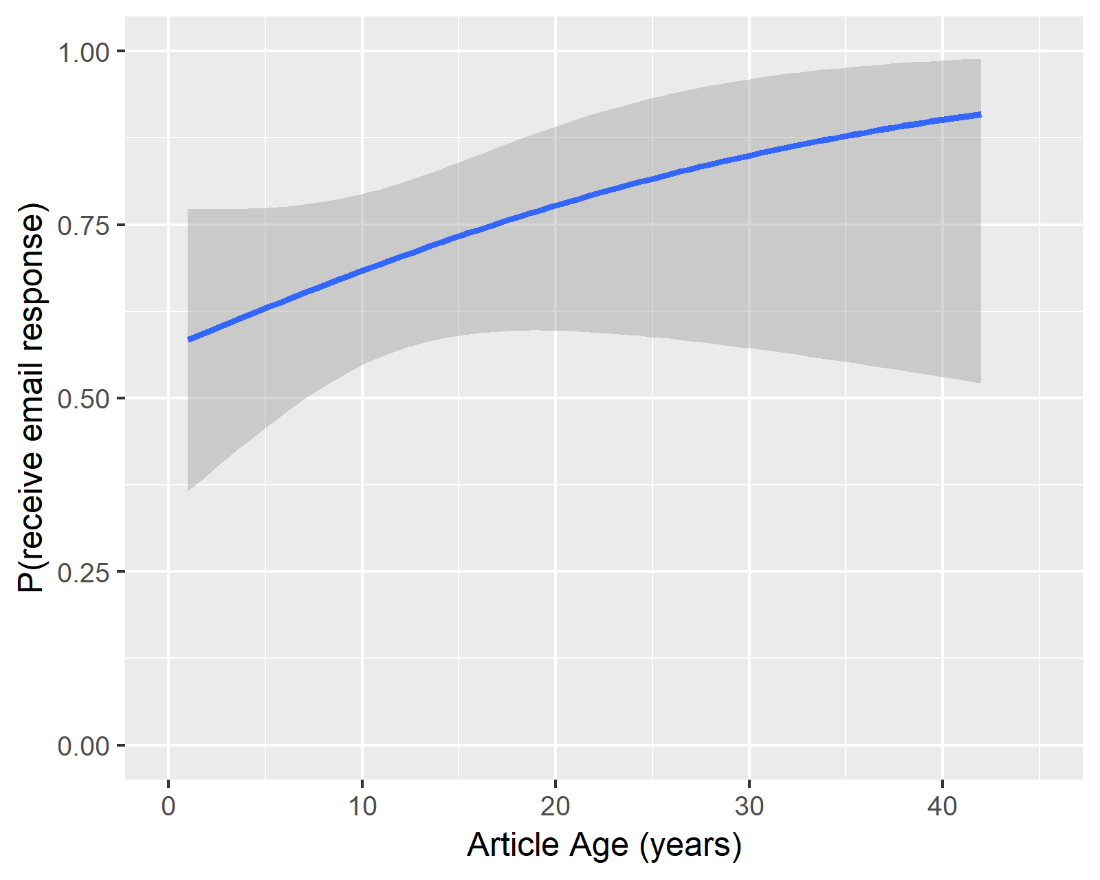

Logistic regression coefficients (se):

Intercept	0.290 (0.480), z-value = 0.605, p = 0.545

Article age (years)	0.0481 (0.035), z-value = 1.370, p = 0.171

The probability to receive response from original authors given a working email could be found does not change significantly over time.

3) Probability that original data were extant (i.e. authors provided us the data, or answered that they were not able to provide us the data due to reasons different from data not extant) given that a working mail could be found and the authors actually responded.

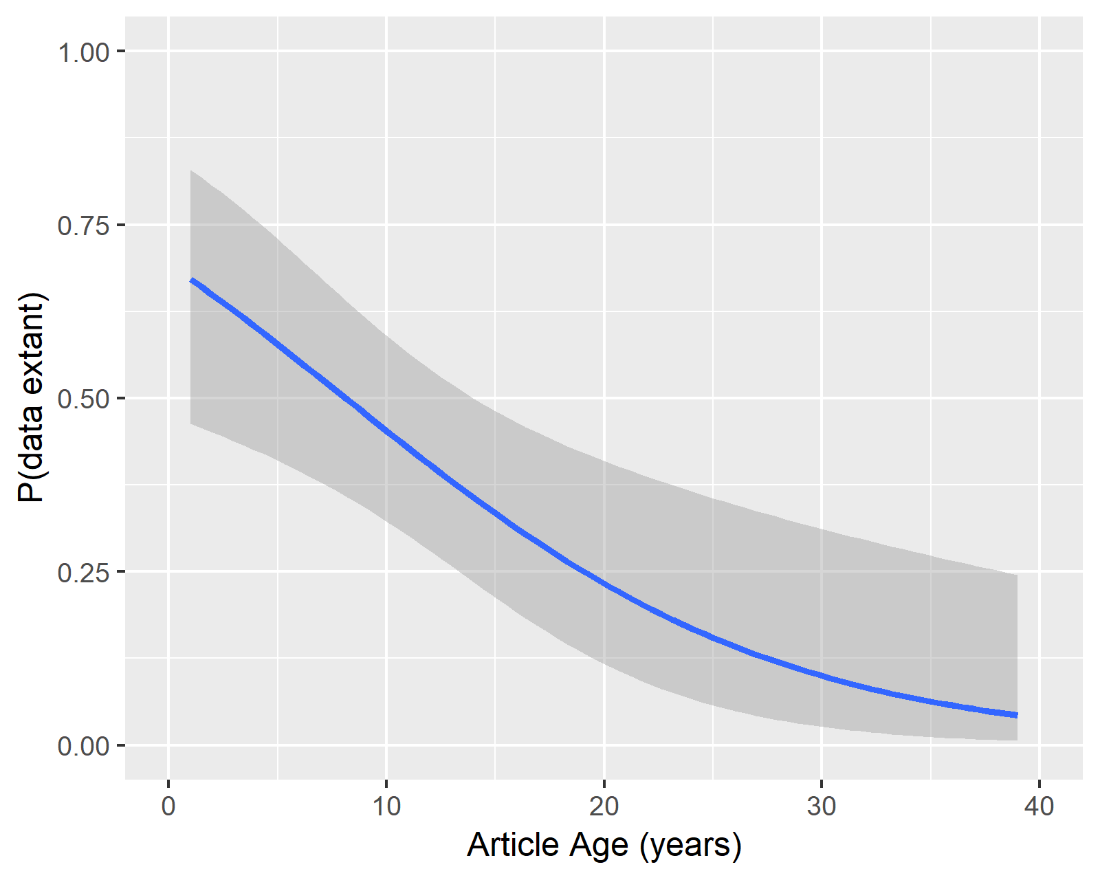

Logistic regression coefficients (se):

Intercept	0.854 ( 0.459), z-value = 1.860, p = .063 .

Article age (years) -0.105 (0.033), z-value = -3.171, p = .00152

The probability that original data were extant declines rapidly with article age.

4) Probability to receive at least some of the requested data given that a working mail could be found and the authors actually responded.

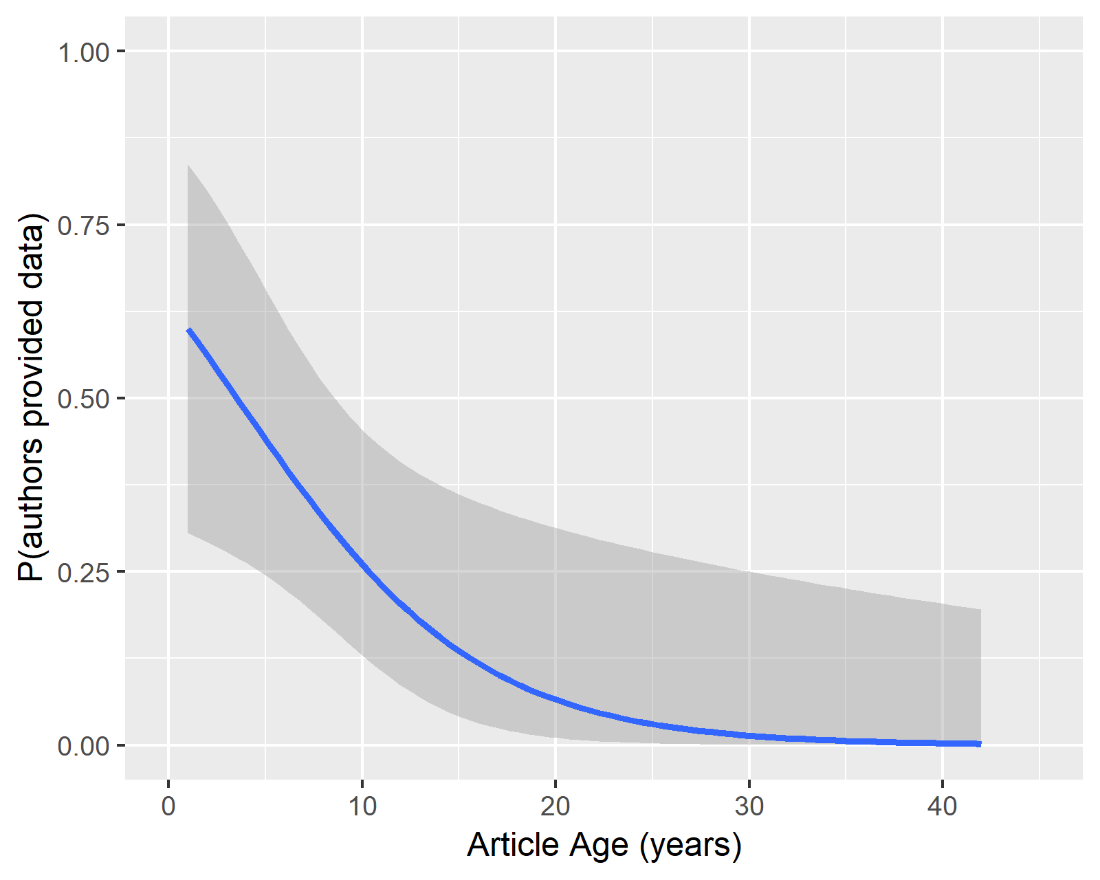

Logistic regression coefficients (se):

Intercept	0.568 (0.682), z-value = 0.833, p = .405.

Article age (years) -0.161 (0.070), z-value = -2.292, p = 0.0219

The probability to receive at least some of the requested data from original authors decreases rapidly with article age. Probability to receive some of the requested data is below 50% only 5 years after article publication, and below 25% only 10 years after article publication.

**Template of the e-mail sent to original authors**

Dear Prof. XXX,

we are working on a systematic review and meta-analysis of acoustic patterns in schizophrenia (for our previous analogous work on autism see: https://www.biorxiv.org/content/early/2016/06/24/046565)

The goal is to provide a more systematic understanding of the cumulative evidence in this area, with a special focus on relation with clinical features of the disorder, and on what should be further investigated. We will be articulating the findings in a report also detailing concerns with sharing data in the field, and how we could work to overcome them in the future. The work is a collaboration between Aarhus University, Aarhus University Hospital and Università di Torino, led by Associate Professor Riccardo Fusaroli (webpage: https://goo.gl/jM9D46).

We are contacting all the authors of the studies included in the review to have a more complete overview of the data, and we considered as eligible for the review your article:

"XXX "

We would be really grateful if you could provide the following:

1. Population-level estimates:

• Mean and standard deviation of the following acoustic measures for groups of individuals with schizophrenia and healthy controls, XXXX.

Without this data we would not be able to include your study in the meta-analysis, so accessing it would be our priority.

2. Additionally, having participant level data from your study would allow us to better assess effects of age, gender and language across studies, as well as possible relations with clinical features. In other words, for each participant we would be really grateful to receive the data in the following list. Note that we are requesting a considerable amount of data and we’d appreciate whatever subset of it you will be able to provide.

• If more than one language is involved in the study, which language is being spoken

• If patients and controls are matched on a one to one basis, some common identifier to account for that in our statistical analysis

• Diagnosis (schizophrenia/control at the least, more specific if available)

• More precise assessment of the clinical features if available (e.g. SANS/SAPS/PANSS scores, general IQ, verbal IQ, etc.)

• If the patients were medicated and the data is available: pharmacotherapy (medication type and dosage)

• Mean and standard deviation of each acoustic feature analyzed (within the participant speech). If different conditions for speech production are employed, we’d be grateful for estimates by participant by condition

• Median and interquartile range of each acoustic feature analyzed if available

• If the participant appears in more than one of your studies, a unique identifier which would allow us to account for the repeated presence of the same participant.

We would also ask you whether you would prefer we keep the data confidential or whether we could share them as an open dataset (as in this example from the previous meta-analysis: https://figshare.com/articles/New_draft_item/3457751).

We know we are requesting a considerable effort on your part, and that you might have ethical and practical concerns with sharing the data, but we really believe this sort of data-sharing will strongly advance our research field.

3. Finally, we are also interested in identifying barriers to open data sharing. If you are not willling/able to share the data with us, it would be very useful if you could share your concerns with us: Why is it problematic to share the data? What could be done to remove this barrier? This will allow us to come up with solutions for future studies.

Many thanks for your work, your time and all your help.

All the best,

Alberto Parola
