## SupplementaryMaterial_RiskOfBias for "Voice Patterns in Schizophrenia: A systematic Review and Bayesian Meta-Analysis"

**Risk of bias**

We used the Cochrane Collaboration tool to assess risk of bias (Higgins and Green, 2011). We identified several potential sources of bias: 1) Matching: the matching criteria between groups of individuals with SCZ and HC differed widely between studies (see Table 1). 2) Clinical heterogeneity: the studies analyzed included samples differing on relevant clinical and cognitive features (e.g. onset, duration of illness, symptoms). In addition, most of the studies may have included more chronic samples of patients with schizophrenia, thus not being fully representative of the entire schizophrenia population. 3) Methodological heterogeneity: the studies analyzed used widely different methods for sample selection, acoustic pre-preprocessing, feature extraction and selection. We extracted and reported in Table 1 all the information that may be relevant for assessing different source of bias. Our aim was to quantify and evaluate the role of these methodological and clinical moderator factors on results of the meta-analysis; however, most of the studies have not reported, or only partially reported this information, and thus there were not enough data points to be analyzed and we were thus not able to quantitatively assess the role of these confounders.

We then measured and tested for heterogeneity of the studies using the Cochran’s Q statistic^13^, which reveals how much of the overall variance can be attributed to true between-study variance, and evaluated publication bias using the rank correlation test (Begg and Mazumdar, 1994). Finally, in the discussion section we tried to synthesize the issues associated with the risk of bias of the included studies, and discussed the impact of heterogeneity we found on results of the meta-analysis.
