## Supplementary material for "Voice Patterns in Schizophrenia: A systematic Review and Bayesian Meta-Analysis": Table_S1_Moderator_analysis_Contrast_SM

**Table S1. Moderator analysis for studies evaluating the effect of diagnosis on acoustic measures**

| **Feature** | **Stacking weight** | **Estimates -Hedges’g**  **95% CI** | | | **Contrast** | **Evidence ratio (credibility)** |
| --- | --- | --- | --- | --- | --- | --- |
|  |  | **Constrained - monologic speech** | **Free - monologic speech** | **Dialogic speech** |  |  |
| Pitch variability | 100% | -0.538 (95% CI = -1.367, 0.238) | -0.111 (95% CI = -2.392, 2.342) | -1.383 (95% CI = -4.49, 1.35) | Free speech > constrained speech | 2.95 (75%) |
|  |  |  |  |  | Dialogic speech < constrained speech | 0.15 (13%) |
| Proportion of spoken time | 100% | -0.385 (95% CI = -2.139, 1.423), | -2.659 (95% CI = -21.195, 6.856) | -1.482 (95% CI = -4.06, 0.832) | Free speech < constrained speech | 0.23 (19%) |
|  |  |  |  |  | Dialogic speech < constrained speech | 4 (98%) |
| Number of pauses | 100% | 0.166 (95% CI =-3.214, 3.241) | -0.092 (95% CI = -4.956, 4.835) | NA | Free speech < constrained speech | 0.72 (42%) |
| Speech rate | 100% | NA | -0.978 (95% CI = -2.423, 0.472) | -0.772 (95% CI = -2.463, 0.862) | Dialogic speech > constrained speech | 1.29 (56%) |
| Intensity variability | 100% | NA | -0.189 (95% CI = -15.307, 14.777) | 1.45 (95% CI = -13.225, 15.363) | Dialogic speech > constrained speech | 1.61 (62%) |
