## Supplementary material for "Voice Patterns in Schizophrenia: A systematic Review and Bayesian Meta-Analysis": Table_S2_Moderator_analysis_Correlations_SM

**Table S2. Moderator analysis for studies evaluating correlations between acoustic measures and clinical symptoms ratings.**

| **Acoustic feature** | **Clinical features** | **Stacking weight** | **Estimates -Hedges’g**  **95% CI** | | | **Contrast** | **Evidence ratio (credibility)** |
| --- | --- | --- | --- | --- | --- | --- | --- |
|  |  |  | **Constrained- monologic speech** | **Free - monologic speech** | **Dialogic speech** |  |  |
| Pitch variability | Positive symptoms | 100% | -0.001 [-24.255, 24.396] | -0.403 [-28.507, 25.236] | 0.79 [-34.599, 37.188] | Free speech < constrained speech | 1.03 (51%) |
|  |  |  |  |  |  | Dialogical speech > constrained speech | 1.11 (53%) |
|  | Negative symptoms | 100% | 0.005 [-1.979, 2.322] | 0.174 [-7.777, 8.697] | -0.431 [-10.764, 7.822] | Free speech > constrained speech | 1.26 (56%) |
|  |  |  |  |  |  | Dialogical speech < constrained speech | 0.87 (46%) |
|  | Alogia | 100% |  | 0.104[-0.504, 0.75] | -0.244 [-0.811, 0.399] | Dialogic speech < free speech | 0.17 (14%) |
|  | Flat affect | 100% |  | 0.026 [-0.264, 0.26] | -0.304 [-0.159, -0.071] | Dialogic speech < free speech | 0.04 (3%) |
| Intensity variability | Flat affect | 100% |  | 0.282 [-2.34, 3.095] | -0.124 [-1.21, 1.218] | Dialogic speech < free speech | 0.34 (25%) |
| Proportion of spoken time | Total psychopathology | 100% | 0.368 [-15.178, 15.608] | - 0.291 [-25.039, 23.229] | -0.152 [-15.01, 14.814] | Free speech < constrained speech | 0.89 (47%) |
|  |  |  |  |  |  | Dialogic speech < constrained speech | 0.78 (44%) |
|  | negative symptoms ratings | 100% | 0.498[95% CI = -7.002, 8.318] | -0.339 [95% CI = -21.721, 20.52] | -0.457 [95% CI = -6.124, 5.166] | Free speech < constrained speech | 0.77 (44%) |
|  |  |  |  |  |  | Dialogic speech < constrained speech | 0.31 (24%) |
|  | Alogia | 100% |  | -0.348 [-8.748, 6.786] | -0.458 [-5.369, 4.584] | Dialogic speech < free speech | 0.71 (41%) |
|  | flat affect | 100% |  | -0.207 [ 95% CI = -3.398, 2.52] | -0.44 [95% CI = -1.507, 0.75] | Dialogic speech < free speech | 0.35 (26%) |
