## Supplementary material for "Voice Patterns in Schizophrenia: A systematic Review and Bayesian Meta-Analysis": Table_S4_Feature_description_SM_060819_RESUB

**Table S4. Description of the acoustic features**

| **Acoustic feature** |  | **Description** |
| --- | --- | --- |
| Fundamental frequency – F0 | Pitch mean | Pitch mean reflects the mean frequency of vibrations of the vocal cords during vocal production across the linguistic unit analysed (phoneme, word, sentence, or the entire speech sample). |
|  | Pitch variability | Pitch variability indicates the mean magnitude of changes in pitch across the linguistic unit analysed (phoneme, word, sentence, or the entire speech sample). |
| Intensity | Intensity mean | Mean intensity or loudness is a measure of the amplitude of the vibrations of the vocal folds, and it represents the mean level of energy carried by a sound wave across the linguistic unit analysed (phoneme, word, sentence, or the entire speech sample). |
|  | Intensity variability | Intensity variability indicates the magnitude of changes in intensity across the linguistic unit analysed (phoneme, word, sentence, or the entire speech sample). |
| Speech production | Duration of utterance | Mean utterance duration indicates the mean duration from beginning to end of a defined speech sample (syllables, word or sentence). |
|  | Speech rate | Speech rate is defined as number of words per time unit (second or minute). |
|  | Percent time talking | Percent time talking represents the percentage of time the speech sample contained a pitch different from zero, i.e. speaking time, relative to the total time of the speech sample. |
|  | Duration of pauses | Mean pause duration indicates the mean duration of pauses across the entire speech sample. |
|  | Number of pauses | Mean number of pauses across the entire speech sample. |
