## Supplementary material for "Voice Patterns in Schizophrenia: A systematic Review and Bayesian Meta-Analysis": Table_S5_Results_meta_analysis_schizotypy_SM

Table S5 - Results schizotypy

| **Features** | **Participants (female) and median** | **Number of studies (articles)** | **Influential study** | **Estimates -Hedges’g**  **[95% CI]** | **P - value** | **ER (Credibility)** | **Sigma squared**  **[95% CI]** | **Q- stats (p-value)** | **Publication bias** |
| --- | --- | --- | --- | --- | --- | --- | --- | --- | --- |
| Pitch Mean | 103 (22 F) SZ  95 (23 F) CT  89 (50 F) ST  26 (14 F) CT | 4(4)  1(1) | No | Effect for SZ vs. ST: -0.07 [-2.522, 2.295]  Effect for ST vs. HC: 0.307 [-1.647, 2.576]  Effect for SZ vs. HC: 0.237 [-0.723, 1.29] | .934  .44  .273 | 1.056 (51%)  2.15 (68%)  3.467 (78%) | 1.199 [0.006, 8.937] | 8.282 (p = .041) | No,  K = 0.2, p = .817 |
| Pitch variability | 387 (92 F) SZ  257 (106 F) CT  75 (56F) ST  100 (68 F) CT | 11(8)  3(3) | No | Effect for SZ vs. ST: -1.577 [-3.076, -0.069]  Effect for ST vs. HC: 1.017 [-0.306, 2.36]  Effect for SZ vs. HC: SZ -0.56 [-1.245, 0.097] | .01  .061  .005 | 45.243 (98%)  13.706 (93%)  99.0 (99%) | 1.294 [0.458, 3.186] | 130.469 (p = .01) | No,  K = -0.297, p = .157 |
| Intensity variability | 104(22 F) SZ  65 (20 F) CT  47 (28 F) ST  73 (41 F) CT | 4 (3)  2 (2) | No | Effect for SZ vs. ST: 1.872 [-2.109, 5.916]  Effect for ST vs. HC: -1.01 [-4.33, 2.227]  Effect for SZ vs. HC: 0.862 [-1.407, 3.306] | .047  .172  .164 | 7.29 (88%)  3.96 (80%)  3.648 (78%) | 5.235 [0.557, 25.367] | 48.939 (p < .001) | No,  K =0.2, p = .719 |
| Proportion of spoken time | 267 (106 F) SZ  211 (98 F) CT  163 (112 F) ST  123 (82 F) CT | 11 (9)  5 (4) | No | Effect for SZ vs. ST: -0.627 [-2.096, 0.813]  Effect for ST vs. HC: - 0.623 [-1.853, 0.53]  Effect for SZ vs. HC: -1.25 [-2.082, -0.428] | .312  .215  .001 | 4.453 (82%)  6.117 (86%)  149.943 (99%) | 1.77 [0.675, 4.098] | 151.321 (p < .001) | No,  K = -0.35, p = .064 |
| Duration of utterance | 93 (30 F) SZ  72 (30 F) CT  75 (56 F) ST  100 (68 F) CT | 4 (4)  3 (3) | No | Effect for SZ vs. ST: 0.924 [-3.358, 4.964]  Effect for ST vs. HC: -1.104 [-4.336, 2.108]  Effect for SZ vs. HC: -0.181 [-2.847, 2.27] | .384  .186  .739 | 2.442 (71%)  4.727 (83%)  1.475 (60%) | 6.903 [1.238, 30.116] | 122.74 (p < .001) | No,  K = -0.429, p = .239 |
| Speech rate | 320 (98F) SZ  243 (99 F) CT  39 (24 F9 ST  37 (23 F) CT | 11 (9)  1 (1) | No | Effect for SZ vs. ST: -2.364 [-5.048, 0.266]  Effect for ST vs. HC: 1.632 [-0.892, 4.263]  Effect for SZ vs. HC: -0.732 [-1.463, 0.033] | .016  .082  .015 | 25.144 (96%)  11.924 (92%)  32.473 (97%) | 1.56 [0.483, 4.466] | 104.414 (p < .001) | No,  K = 0.0, p = 1.0 |
| Duration of pauses | 221 (128 F) SZ  150 (92F) CT  39 (24 F) ST  37 (23F) CT | 9 (8)  1 (1) | No | Effect for SZ vs. ST: 3.402 [-0.392, 7.276]  Effect for ST vs. HC: -1.541 [-5.299, 2.066]  Effect for SZ vs. HC: 1.861 [0.719, 3.055] | .009  .212  < .001 | 27.881 (97%)  4.844 (83%)  234.29 (100%) | 3.039 [0.704, 9.806] | 75.624 (p <. 001) | Yes,  K = 0.6 p = .017 |
| Number of pauses | 68 (23 F) SZ  40 (13 F) CT  47 (28 F) ST  73 (41 F) CT | 5 (4)  2 (2) | No | Effect for SZ vs. ST: -1.42 [-4.191, 1.356]  Effect for ST vs. HC: 1.398 [-1.086, 3.778]  Effect for SZ vs. HC: -0.021 [-1.617, 1.478] | .078  .033  .782 | 7.715 (89%)  9.753 (91%)  1.321 (57%) | 2.814 [0.381, 13.076] | 39.83 (p < .001) | No,  K = -0.619, p = .069 |
