## Supplementary material for "Voice Patterns in Schizophrenia: A systematic Review and Bayesian Meta-Analysis": Table_S3_Machine_Learning_SM

**Table S3. Multivariate machine learning studies**

| **Study** | **Aim** | **Classifier** | **Cross-validation** | **Acoustic features^[1]^** | **Index and results** |
| --- | --- | --- | --- | --- | --- |
| Rapcan et al. (2009)^[2]^ | Predict diagnosis | LDA | Cross-fold validation | Number of Pauses, Proportion of Silence, Total Recording Time, Total Length of Pauses (s). | Sensitivity (%) 72.64 75.21 72.36  Specificity (%) 78.63 83.62 85.47  Positive predictivity (%) 77.86 82.02 83.39  Negative predictivity (%) 74.24 77.40 75.75  Overall accuracy (%) 75.64 79.42 78.92  Area under the ROC curve 0.79 0.82 0.80 |
| Martinez-Sanchez et al. (2015) | Predict diagnosis | LDA | NR | Task duration (s), Intensity (dB), Pause rate, Pitch Mean F0 (Hz), Pitch variability SD (Hz), F0 Range (ST), Syllabic dynamics, Prosodic peaks, Prosodic valleys, Intra-syllabic Trajectory (ST/s), Inter- syllabic Trajectory (ST/s), Phonation Trajectory (ST/s) | Overall accuracy: 87.5% |
| Kliper et al. (2015) | Predict diagnosis | SVM | Leave-one-out | Spoken Ratio, Utterance Duration, Gap Duration, Pitch Range, Pitch variability, Power variability, Mean Waveform Correlation (MWC), Jitter, Shimmer | Overall accuracy: 76.19% |
| Puschel et al. (1998) | Predict diagnosis | LDA | NA | Total recording time, total length of utterances, number of pauses, mean energy per second, variation of energy per second and JO-contour. | Overall accuracy: 85.6%  3.6 % false positive  22.6 % false negative  After 2 weeks:  overall accuracy: 83.3%.  4.4 % false positive  28.9 % false negative |
| Puschel et al. (1998) | Predict symptoms | LDA | NA | Total recording time, total length of utterances, number of pauses, mean energy per second, variation of energy per second and JO-contour. | Overall accuracy: 78.6%  After 2 weeks: 71.4%. |
| Stassen et al. (1995) | Predict diagnosis | LDA | NA | Mean utterance duration, mean energy per second, variation of energy per second, mean pause duration per second, mean vocal pitch, F0 – amplitude, F0 - contour | Overall accuracy: > 80%. |
| Stassen et al. (1995) | Predict symptoms | LDA | NA | Mean utterance duration, variance of utterance duration, mean energy per second, variation of energy per second, mean vocal pitch, F0 – amplitude, F0 – bandwidth, F0 - contour | Overall accuracy for negative symptoms: 75.9%  Overall accuracy for positive symptoms: 71.9% |

LDA: Linear Discriminant Analysis; SVM: Support-vector machines

1. Set of acoustic features used to train the ML classifier
2. The authors used three different features sets to train the classifier, we reported results for each features set. For additional details see Rapcan et al. (2009), page4.
